# Monkeys learn to report their own sensory cortical population activity

**DOI:** 10.64898/2026.08.07.743408

**Authors:** Jiaqi Hu, Gouki Okazawa

## Abstract

How sensory cortical activity is read out by downstream circuits to guide behavior is a fundamental unsolved problem. Substantial work has examined correlations between sensory neural responses and animals’ perceptual judgments, but their interpretations remained controversial due to many intervening variables, such as responses of unrecorded neurons. Furthermore, stimulus and choice encoding in sensory populations are often not well aligned, and it remains contested whether this misalignment indicates a limitation in sensory readout. Here, we introduce a closed-loop, neurofeedback paradigm that directly interrogates the capacity of sensory readout: the key idea is to train subjects to report specific patterns of population activity in a sensory area recorded online, rather than the actual stimuli presented. To test this, we trained macaque monkeys on a visual change-detection task using shape stimuli, implanted an electrode array in visual area V4, and tested whether they could be further trained to rely on their own V4 activity along specific axes in neural state space. Strikingly, monkeys successfully increased the neuron-choice correlation along trained axes in neural population state space. No detectable changes in stimulus selectivity or noise correlations were found within the recorded population, and further model simulations confirmed that adjustment of sensory readouts best accounted for the results. A control experiment that merely disrupted the stimulus–reward contingency without a closed loop failed to enhance neuron–choice correlation. Together, these results demonstrate that closed-loop neural feedback achieves neuron-choice alignment beyond the ceiling of natural perceptual training, suggesting that the misalignment in perceptual tasks reflects constraints on learning within naturally available training regimes.

## Introduction

How neural activity in the nervous system gives rise to perception has long been a central question in systems neuroscience. For decades, researchers have addressed this question by training animals to make perceptual judgments on ambiguous (near-threshold) sensory stimuli and probing the correlation between their behavioral reports and the activity of individual neurons (Britten et al. 1996; Pletenev et al. 2026). Positive neuron-choice correlations (often quantified as “choice probability”; Britten et al. 1996) have been consistently observed across various sensory cortices, and earlier works considered it as evidence for a link between neural activity and perception (Parker and Newsome 1998). However, many other interpretations have been proposed since then; theories have shown that choice probability strongly depends on the correlation structure within the neural population (Cohen and Newsome 2009; Cumming and Nienborg 2016; Nienborg and Cumming 2010; Shadlen et al. 1996) rather than on the direct contribution of the recorded neuron to perceptual reports (Haefner et al. 2013). Non-causal correlations can also arise from feedback of choice-related signals from downstream areas to sensory cortices (Zhao et al. 2020). Empirically, studies have revealed that choice probability deviated from other measures that assessed the effects of stimuli or sensory neurons on behavior, such as microstimulation (Yu and Gu 2018) or psychophysical reverse correlation (Levi et al. 2023; Nienborg and Cumming 2009).

More recently, the proliferation of large-scale neural recordings has extended this question to the population level, but the interpretation of neuron-choice correlations remains controversial. Analyses of simultaneously recorded population activity in sensory areas have revealed axes in neural state space that can decode animals’ choices. However, these choice axes are often misaligned with the axes that best decode the stimuli the animals are trained to discriminate (Ni et al. 2018, 2022; Pletenev et al. 2026; Weiner and Ghose 2014; Zhao et al. 2020). The misalignment suggests suboptimal information use, but its origin remains unresolved. One possibility again is top-down feedback of choice-related signals into a subspace orthogonal to stimuli (Zhao et al. 2020), but choice-related activity is also observed in early sensory responses before feedback would arrive (Britten et al. 1996; Wimmer et al. 2015). Some have proposed that the misalignment reflects an ecological strategy of animals to detect diverse stimuli (Ni et al. 2022; Weiner and Ghose 2014) or to efficiently learn tasks (Laamerad et al. 2025). It is also noted that this choice axis tends to align with the axis of greatest variance in neural state space (Ni et al. 2018), leading to the idea that it may be selected for effective information transmission (Haimerl et al. 2023; Nassar et al. 2021; Panzeri et al. 2022; Srinath et al. 2025).

A critical limitation that has hindered deeper investigation is that typical perceptual tasks can only indirectly link sensory neural activity and animals’ choices; sensory inputs inevitably activate widespread brain networks that contribute to behavior, yet experiments can only measure correlations between behavior and a small subset of recorded sensory neurons, leaving many unobservable variables unaccounted for. If the goal is to show whether recorded neurons influence perception and behavioral choices, a direct approach would be to manipulate their activity using microstimulation or other causal interventions. However, the limited precision of these techniques constrains our ability to address more computationally motivated questions (Jazayeri and Afraz 2017), such as what axes in neural state space guide behavior (Semedo et al. 2019).

Here, we introduce a closed-loop paradigm that directly interrogates the capacity of sensory readout; rather than training animals to perform sensory judgments, we train them to report specific patterns of sensory population activity recorded online, directly linking behavioral feedback to neural signals in a population subspace. This approach narrows the gap between behavioral choices and recorded neural signals in a way that typical perceptual tasks cannot and goes beyond coarse perturbation methods by precisely targeting specific patterns of population activity. We first trained monkeys on a visual change-detection task using shape stimuli (Fig. 1A, left), implanted an electrode array in visual area V4, and then tested whether closed-loop training (Fig. 1A, right) could align their choices with V4 activity along axes encoding stimulus features in neural state space. Monkeys successfully increased neuron-choice correlation along these axes without any detectable change in V4 response properties. The increased correlations suggest that the closed-loop training can achieve neuron-choice alignment beyond the ceiling of natural perceptual training.

**Figure 1:**
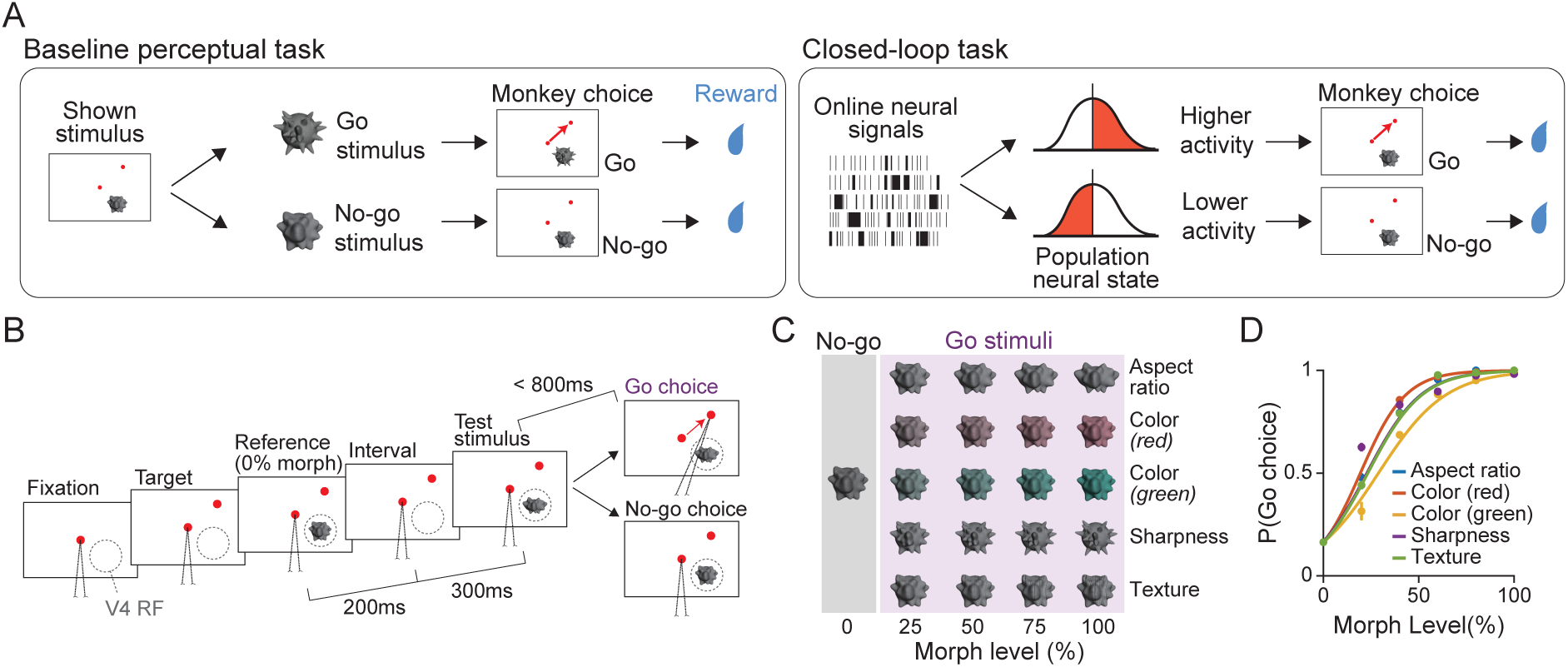
Baseline change detection task and closed-loop paradigm. (**A**) In standard perceptual tasks, subjects make a choice based on physical stimuli (left). In the proposed closed-loop task (right), neural population activity from a sensory area is recorded during stimulus presentation, and the correct choice is determined by the online neural activity; for example, if activity along a specific axis in neural state space exceeds a threshold, a Go response becomes correct. (**B**) Trial structure for the baseline change-detection task. After fixation and the appearance of a peripheral target dot, a reference and a test stimulus were presented sequentially with a brief inter-stimulus interval. Monkeys were required to detect any change in the test stimulus relative to the reference and, if detected, make a saccade to the target, or otherwise maintain fixation for 800 ms. (**C**) Object features used in the task (each row). We varied morph levels to adjust stimulus difficulty. The reference was always the 0% morph. Test stimuli with positive morphs (∼ 50% of trials) required a Go response for reward. During training, one feature was selected randomly on each trial. During the main experiments, a single feature was used throughout each session to enable comparison between baseline and closed-loop conditions. (**D**) Psychometric functions in the change-detection task. Example data from monkey B. Error bars indicate S.E.M. across sessions. See Supplementary Fig. 1 for monkey A.

## Results

### Stimulus and choice encoding are misaligned in the change-detection task

We first extensively trained two macaque monkeys on a change-detection task, in which they were required to detect the changes in the shape, color, and texture of sequentially presented object stimuli (Fig. 1B). After monkeys fixated on a central dot, a peripheral target dot appeared, followed by reference and test stimuli presented with a brief inter-stimulus interval at the average receptive field (RF) location of the recorded V4 population. Monkeys were required to make a saccade to the target upon detecting any difference between the reference and test stimulus, and to maintain fixation when no change was detected. The saccade target was positioned approximately orthogonal to the V4 RF to minimize contamination from saccade-related signals in V4 (Chandrasekaran et al. 2024; Steinmetz and Moore 2014). On each trial, one of 4-5 features (color, texture, sharpness, aspect ratio; Fig. 1C) could change, with a change magnitude defined as a morph level ranging from 0% (no change) to 100% (full change), scaled based on preliminary human psychophysics (see Methods). The reference stimulus was always at 0% morph. Thus, when the test stimulus was not 0% morph (shown in *∼* 50% of trials), choosing a Go target yielded a reward. Both monkeys successfully learned the detection task (Fig. 1D), achieving performance comparable to, or slightly below, that of humans (Supplementary Fig. 1). This change-detection training set the baseline for the neuron-choice correlations obtained in natural perceptual tasks and also prepared them for the forthcoming closed-loop experiments. In the main experiments explained below, only one feature was varied per session.

The V4 population, recorded with a chronically implanted 96-channel Utah array, encoded the strength of each stimulus feature along an approximately linear axis within population state space. Many units showed significant selectivity for the varied features (57.2%, 83.7% in monkey A and B, respectively; one-way ANOVA), and most selective units showed higher firing rates at higher morph levels (Fig. 2A; average of selective units). To minimize the influence of feedback from downstream areas, subsequent analyses and closed-loop experiments were restricted to an earlier stimulus presentation period (80 to 150 ms from stimulus onset; gray shade in Fig. 2A). We calculated mean firing rates for each trial and unit and denoised them by reducing the population dimensionality to 12 via principal component analysis (PCA). The 12 dimensions were chosen through a parallel analysis (Altan et al. 2021; Supplementary Fig. 2A). We then applied linear regression to predict the stimulus morph level on each trial. Mean population responses to different morph levels fell approximately along a line in the population state space (Fig. 2B), confirmed by the high accuracy of linear regression fit to the mean responses at each morph level (Fig. 2C).

**Figure 2:**
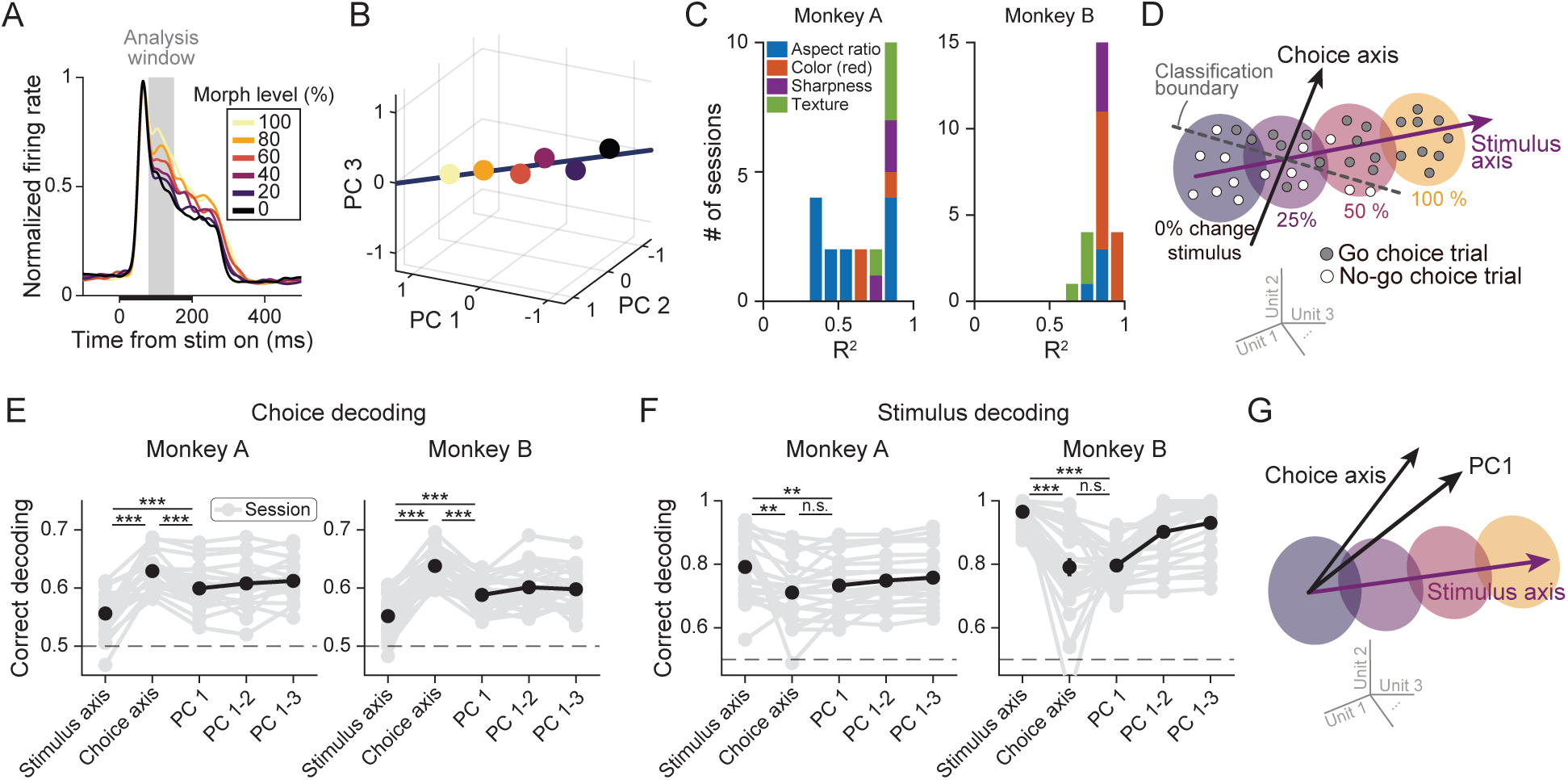
Misalignment of stimulus and choice encoding axes in V4 during the change-detection task. (**A**) Example population average responses to object stimuli (red color morph). The shaded region indicates the time window used for all analyses and closed-loop experiments. The example is from a passive-fixation session from monkey A (black horizontal bar: stimulus period), but the response properties were similar in the main tasks. (**B**) Average population responses to stimuli with different morph levels lie approximately along a linear axis in neural state space, visualized using PCA for the same session with A. Circles represent trial-averaged population response to each morph level. (**C**) A linear regression of neural population responses onto stimulus morph levels explained substantial variance in the 12D latent PC space. Colors indicate the object feature presented in that session. (**D**) Definitions of stimulus and choice axes in neural state space. As demonstrated in **C**, the stimulus axis can be defined as a linear axis in the space. The choice axis is defined as the axis that best separates Go and No-go choices after removing the effect of stimulus morph levels. (**E**) Choice decoding accuracy was significantly lower along the stimulus axis than along the choice axis, indicating their misalignment. The dominant axes of population variability (top PCs) decode choice more accurately than the stimulus axis. A dashed horizontal line indicates the chance level. Black circles represent session means, and error bars indicate S.E.M. across sessions (not visible due to small values). (**F**) Conversely, stimulus decoding accuracy (0% versus positive morphs) was significantly lower along the choice axis than along the stimulus axis, confirming their misalignment. The dominant axes of population variability (top PCs) also carried less stimulus information than the stimulus axis. (**G**) Schematic. The stimulus and choice axes are misaligned, and the choice axis is more closely aligned with the dominant axis of population variability (PC1).* p < 0.05, ** p < 0.01., *** p < 0.001.

How was stimulus-selective population activity related to monkey choice? Although Go choices must generally be associated with activity along the positive-morph side of the stimulus encoding axis, the neural activity predictive of monkey choice may not be fully aligned with stimulus encoding (Fig. 2D; Ni et al. 2018, 2022; Weiner and Ghose 2014; Zhao et al. 2020). To test this, we identified the choice decoding axis by subtracting the mean activity for each morph level along each dimension of the latent space and then performing a linear logistic regression that classified Go and No-go choices. Choice decoding performance along the obtained axis (“choice axis” in Fig. 2E) —evaluated with cross validation— was comparable to previous V4 studies (Jasper et al. 2019; Ni et al. 2018; Mean decoding accuracy: 0.629 for monkey A, *t*(21) = 20.15, *p <* 10*^−^*^10^; 0.638 for monkey B, *t*(27) = 27.35, *p <* 10*^−^*^10^, two-tailed *t*-test). However, choice decoding accuracy along the stimulus axis (“stimulus axis” in Fig. 2E) was substantially lower than that along the choice axis (0.556 for monkey A, *t*(21) = *−*10.13, *p* = 1.5 *×* 10*^−^*^9^; 0.552 for monkey B, *t*(27) = *−*12.65, *p <* 10*^−^*^10^) albeit being above chance (*t*(21) = 7.90, *p* = 1.0 *×* 10*^−^*^7^ for monkey A; *t*(27) = 8.54, *p* = 3.7 *×* 10*^−^*^9^ for monkey B). Conversely, stimulus decoding performance (0 % vs. non 0% morph stimuli) along the choice axis was poorer than that along the stimulus axis (Fig. 2F; *t*(21) = 3.27, *p* = 0.0037 for monkey A and *t*(27) = 6.36, *p* = 8.3 *×* 10*^−^*^7^ for monkey B, two-tailed *t*-test). Together, these results indicate that stimulus and choice information were not fully aligned in population state space (Fig. 2G), consistent with previous observations (Ni et al. 2018, 2022; Pletenev et al. 2026; Weiner and Ghose 2014; Zhao et al. 2020).

We further asked whether intrinsic neural fluctuations were aligned with the choice axis (Haimerl et al. 2023; Nassar et al. 2021; Panzeri et al. 2022; Srinath et al. 2025). We extracted the leading PCs of the 12-dimensional subspace as axes capturing major fluctuations in population activity. This PCA was performed on neural responses during the stimulus period (gray shading in

Fig. 2A), but PC scores remained largely stable across wider trial epochs (Supplementary Fig. 2B, C). The first PC decoded monkey choice better than the stimulus axis (Fig. 2E; *t*(21) = *−*4.84, *p* = 8.8 *×* 10*^−^*^5^ for monkey A and *t*(27) = *−*5.21, *p* = 1.7 *×* 10*^−^*^5^ for monkey B, two-tailed *t*-test), while its decoding performance for stimulus was indistinguishable from that of the choice axis (Fig. 2F; *t*(21) = *−*1.98, *p* = 0.061 for monkey A and *t*(27) = *−*0.15, *p* = 0.87 for monkey B, two-tailed *t*-test). Thus, the choice axis was misaligned with the stimulus encoding axis, while it was closer to the dominant axis of intrinsic neural fluctuations (Fig. 2G). These results are also consistent with previous works (Ni et al. 2018; Srinath et al. 2025), while the source of this stimulus-choice misalignment remains contested (Laamerad et al. 2025; Ni et al. 2022).

### Closed-loop training increased neuron-choice correlation

We then conducted the main closed-loop experimental sessions (Fig. 3A). In this experiment, we projected the recorded V4 population activity onto a predefined stimulus axis in real time and trained the monkey to align their choices with the decoder output rather than with the physical stimulus. The stimulus decoding axis was determined during a passive viewing task performed immediately before each closed-loop session. During the closed-loop experiments, the correct answer on each trial was set by comparing V4 responses along the stimulus axis to a fixed threshold (Fig. 3A inset). To quantify closed-loop training effects, we first established a behavioral baseline on the change-detection task using a single object feature over 4 or more sessions, then conducted 5-7 closed-loop sessions with the corresponding stimulus axis as a training target (Fig. 3B). This procedure was then repeated for a different object feature in the next round.

**Figure 3:**
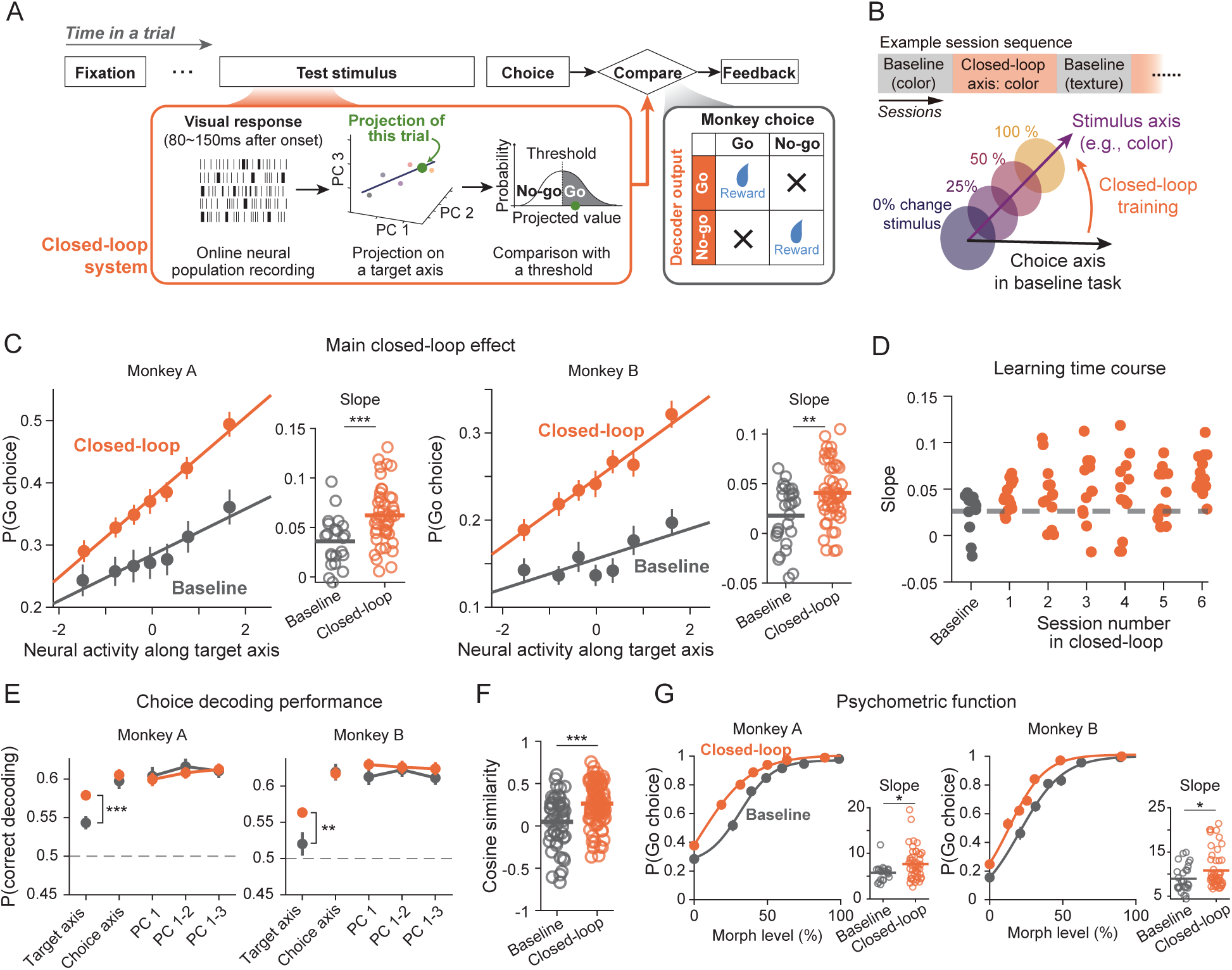
Closed-loop training improved neuron-choice correlation. (**A**) During the closed-loop task, neural population responses were analyzed in real time to determine the correct choice. Spike counts were projected onto a predefined stimulus axis in the latent state space, and if the projected value exceeded a threshold, a Go response became correct (orange and gray inset). (**B**) (top) In each round of data collection, monkeys performed the standard change-detection task with a single object feature as a baseline (≥ 4 sessions), followed by the closed-loop task with the same object feature (≥ 6 sessions). (bottom) We set the stimulus encoding axis as the closed-loop target. (**C**) Both monkeys improved neuron-choice correlation through closed-loop training. (left) Go response probability calculated for binned neural activity along the target stimulus axis, averaged across sessions. Error bars indicate S.E.M. (right) Slopes estimated from individual sessions (open circles). Horizontal bars indicate the means. (**D**) Slopes over sessions. Baseline slopes were calculated by pooling all baseline sessions within each round. The gray horizontal line is the average of the baseline. Orange dots are individual sessions. (**E**) In both monkeys, choice decoding accuracy along the stimulus axis increased, while the accuracy along the choice axis and the PC1 remained unchanged. (**F**) Cosine similarity between the target stimulus axis and choice axis increased after closed-loop training. (**G**) The slopes of psychometric functions (α_1_ in Eq. 3) were slightly steeper after closed-loop training in both monkeys. * p < 0.05, ** p < 0.01, *** p < 0.001.

Strikingly, closed-loop training induced a robust increase in neuron-choice correlation. To quantify this, we plotted the probability of Go choice as a function of neural activity projected onto the trained stimulus axis (Fig. 3C; binned into seven levels), focusing on 0% morph trials in which no physical stimulus change occurred. The slope of this function reflects the strength of correlation between monkey choice and projected neural activity. In the baseline sessions, the slopes were already significantly greater than zero for both monkeys (Fig. 3C, gray; *t*(21) = 6.75, *p* = 1.1 *×* 10*^−^*^6^ for monkey A, *t*(27) = 3.11, *p* = 0.004 for monkey B, two-tailed *t*-test), consistent with the above-chance choice decoding along the stimulus axis observed earlier (Fig. 2E). Critically, the slopes became significantly steeper following closed-loop training in both animals (Fig. 3C; *t*(66) = 3.66, *p* = 5.1 *×* 10*^−^*^4^ for monkey A and *t*(76) = 3.23, *p* = 0.002 for monkey B, two-tailed *t*-test across sessions), indicating enhanced correlation between population activity and choice.

This effect remained evident when non-zero morph trials were included (Supplementary Fig. 3A). Furthermore, choice probability calculated for individual units exhibited greater correlations with their weights for the closed-loop axis after the training (Supplementary Fig. 3B). When we plotted the time course across sessions within each round, we found an increasing trend from day 1 of the closed-loop training, which persisted over sessions (Fig. 3D).

The steeper neuron-choice slopes reflected a better alignment between the stimulus and choice axes, rather than an increase in overall choice information in V4. As in the previous section, we compared choice decoding accuracy along the stimulus axis against that along the best choice decoding axis. Consistent with the change in slopes, choice information captured along the stimulus axis increased significantly after closed-loop training (Fig. 3E; *t*(66) = 3.69, *p* = 4.5 *×* 10*^−^*^4^ for monkey A and *t*(76) = 2.87, *p* = 0.005 for monkey B, two-tailed *t*-test). Importantly, however, the overall magnitude of choice information remained unchanged, as evidenced by stable choice decoding accuracy along both the choice axis (*t*(66) = 0.61, *p* = 0.55 for monkey A and *t*(76) = *−*0.12, *p* = 0.91 for monkey B, two-tailed *t*-test) and the leading PC axes (PC1: *t*(66) = 0.61, *p* = 0.76 for monkey A and *t*(76) = 1.25, *p* = 0.22 for monkey B). We further quantified axis alignment between the choice and stimulus axes through their cosine similarity (Fig. 3F), which was significantly greater after closed-loop training than at baseline (*t*(144) = 4.60, *p* = 9.1 *×* 10*^−^*^6^, two-tailed *t*-test). Together, closed-loop training strengthened the neuron-choice correlation by aligning the stimulus and choice axes without altering the total amount of choice-related activity in V4.

Did closed-loop training also alter behavioral performance on the physical change detection? Interestingly, we found both monkeys showed a moderate increase in psychometric slope during closed-loop training (Fig. 3G; *t*(66) = 2.43, *p* = 0.018 for monkey A and *t*(76) = 2.11, *p* = 0.038 for monkey B, two-tailed *t*-test), indicating improved perceptual sensitivity. This could be an outcome of the better alignment between the stimulus and choice axes. One might wonder whether heightened attention or engagement during closed-loop sessions could have independently improved both psychometric slope and neuron-choice alignment, but analyses in the next section weigh against this possibility.

We also observed an increase in overall Go response probability following closed-loop training, evident in both neuron-choice slopes (Fig. 3C) and psychometric functions (Fig. 3G). This was not an intended outcome; we speculate that the broken contingency between stimuli and correct choices (e.g., 0% morph trials could be rewarded by choosing Go) encouraged the monkeys to bias their choices. There was no significant correlation between this Go bias and neuron-choice correlation across sessions (*p >* 0.27 for both monkeys; Supplementary Fig. 4), ruling out the possibility that this bias somehow caused the increased neuron-choice correlation. Also, we later present a control experiment showing that the broken contingency alone could not drive the neuron-choice correlation.

### Changes in neural selectivity and correlation structures do not explain the closed-loop effects

Since choice probability can be strongly influenced by neurons’ stimulus selectivity (Pitkow et al. 2015) and pairwise-correlation structures among neural populations (Haefner et al. 2013), we wondered whether changes in these properties underlie closed-loop effects. For example, heightened attention or perceptual learning can sharpen neural tuning (Reynolds et al. 2000; Yang and Maunsell 2004) or reduce noise correlations (Cohen and Maunsell 2009; Mitchell et al. 2009), either of which could enhance neuron-choice correlations (Law and Gold 2009). We therefore tested whether closed-loop training produced any systematic changes in neural response properties.

We first assessed whether stimulus encoding improved during closed-loop training and found no substantial evidence for it. We measured classification accuracy between the 0% morph and positive morph stimuli at each morph level along the best stimulus decoding axis in the 12-dimensional latent space. The resulting neurometric functions increased monotonically with morph level but their slopes were indistinguishable between baseline and closed-loop conditions (Fig. 4A; *t*(144) = 0.18, *p* = 0.86, two-tailed *t*-test, sessions aggregated across monkeys; for individual monkeys, see Supplementary Fig. 5). Likewise, response variability, quantified as the Fano factor computed on 0% morph trials, did not significantly differ (Fig. 4B; *t*(144) = 0.37, *p* = 0.71, two-tailed *t*-test).

**Figure 4:**
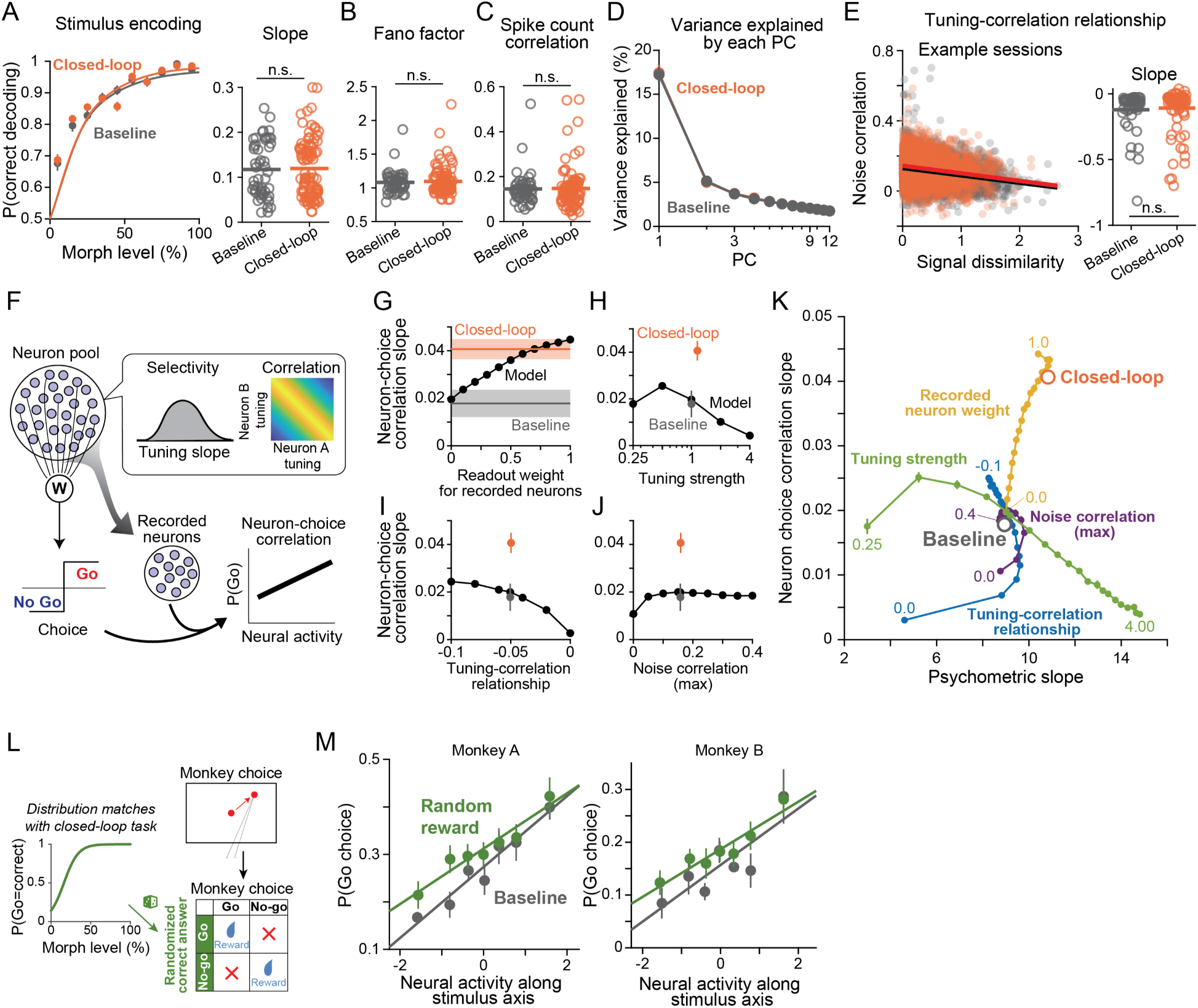
Neuronal response properties did not change after closed-loop training, supporting a readout account. (**A** to **E**) Stimulus encoding and correlation structure remained unchanged after closed-loop training. We assessed population stimulus decoding accuracy (positive versus 0% morph; **A** left) quantified as neurometric slopes (**A**, right), Fano factor (**B**), pairwise spike count correlations (**C**), population correlation structure quantified as eigenvalues of principal components (**D**), and the relationship between stimulus tuning similarity and noise correlation (**E**). Open circles are individual sessions. Pale dots in **E** are individual unit pairs. Results are pooled across monkeys. See Supplementary Fig. 5 for individual monkey results. (**F**) A model consisting of sensory neuron pools (n = 10, 000) whose selectivity and noise correlation structure were matched to the recorded data. A linear weighted sum of the activity determined a Go/No-go choice. Neuron-choice correlation was calculated from a subset of neurons (n = 100) to simulate realistic recording conditions. (**G**) Aligning the readout weights with the recorded population increased neuron-choice correlation to a level comparable to that observed in the monkey data. Shading on the lines indicates S.E.M. of the empirical data from monkey B. Monkey A’s data could also be reproduced. (**H** to **J**) Modifying response properties of the neuron pool alters neuron–choice correlation, but the magnitude of change required to match the closed-loop data fell outside the range of response property changes observed in the empirical data, ruling out these alternatives. (**K**) Diagram illustrating how each model parameter affects psychometric slope (behavioral response) and neuron-choice correlation slope. Gray and orange open circles represent experimental data. Each line corresponds to independent simulations varying a single parameter. Numbers adjacent to lines indicate the parameter value. Error bars on model results indicate S.D. across 5 model instances. (**L**) We also performed a random reward control task to confirm that the neuron–choice correlation does not increase without closed-loop. In this task, stimulus-target contingency was disrupted as in the closed-loop task, but reward delivery was determined randomly rather than by online neural activity. On each trial, a Go response was rewarded with a probability matched to that of the trials with the corresponding morph level in the closed-loop sessions of the same monkey. The probability shown in the inset is an example from monkey B. (**M**) Neither monkey improved neuron–choice correlation along the stimulus axis in this task.

We next assessed pairwise noise correlations and again found no substantial change following closed-loop training. Mean spike count correlation across all recorded channels, computed on 0% morph trials, did not differ significantly (Fig. 4C; *t*(144) = 0.17, *p* = 0.87, two-tailed *t*-test). We further characterized the structure of pairwise correlations by computing the eigenvalue spectrum of the covariance matrix across all recorded units. No significant change after training was observed (Fig. 4D; PC1: *t*(144) = 0.15, *p* = 0.88, two-tailed *t*-test). We also examined the relationship between neuronal tuning and pairwise correlation structure (Haefner et al. 2013). Each neuron was approximately linearly tuned to morph levels (Fig. 2A-C), thus we used a linear regression slope as a measure of tuning strength. Consistent with prior reports (Kohn and Smith 2005), neuron pairs with similar tuning tended to exhibit higher spike count correlations (Fig. 4E left; baseline example session: *r* = *−*0.25, *p <* 10*^−^*^10^). However, the strength of this relationship was unchanged following closed-loop training (Fig. 4E right; *t*(144) = 0.47, *p* = 0.64, two-tailed *t*-test). These results held consistently across both monkeys (Supplementary Fig. 5).

Given the absence of detectable changes in V4 response properties, we developed a neural population model to test whether changes in behavioral readout alone could account for the closed-loop effects (Fig. 4F). The model consisted of a pool of 10,000 simulated V4 units whose stimulus tuning, pairwise correlations, and the tuning-correlation relationship were approximately matched to parameters sampled from the recorded data. The model linearly sums unit firing rates according to readout weights and generates a binary Go/No-go choice by comparing the result to a fixed threshold. To mimic the electrophysiological recordings, we randomly subsampled 100 units from the pool and computed neuron–choice correlation.

We first confirmed that this population model reproduced monkeys’ psychometric curves and neuron-choice correlation slopes in the baseline condition (Supplementary Fig. 6B). Neural response properties were held fixed, while several free parameters, such as the decision threshold, were adjusted to match these experimental data (see Methods). Readout weights were constructed by adding random weights to the optimal stimulus decoder of the whole unit pool (Pitkow et al. 2015), mimicking the misalignment of stimulus and choice axes. We then held all of them fixed except for the parameter under investigation to test which parameters successfully account for the closed-loop effects.

Changes in readout weights could reproduce the main results. Increasing the readout weights for the 100 sub-sampled units enhanced the neuron-choice correlation (Fig. 4G) with a slight improvement in the psychometric curves (Fig. 4K, yellow line), matching the experimental data in the closed-loop sessions. Notably, to match the data, readout weights had to be shifted substantially toward the recorded units (Fig. 4G; *w* = 1 corresponds to full weighting on the recorded subpopulation), suggesting that the effect sizes observed experimentally could have been approaching a theoretical limit of possible correlations.

By contrast, modifying response properties of the units could not reproduce the results within meaningful parameter ranges. Parameters such as individual neurons’ tuning slope (Fig. 4H) or the relationship between tuning dissimilarity and noise correlation (Fig. 4I) could influence the neuron-choice correlation, but the magnitude of changes required to approach the data pushed these parameters far outside the range observed experimentally (Fig. 4H-J). Changes in decision threshold (i.e., Go choice rate) likewise failed to produce a comparable slope increase (Supplementary Fig. 6C). Lastly, the joint visualization of the neuron–choice correlation and psychometric slopes confirmed that, of the parameters tested, only readout weight changes reproduced the closed-loop effects (Fig. 4K).

### The broken stimulus-target contingency does not account for the effects

Our closed-loop design linked internal neural activity to correct targets, but, as a consequence, partially broke the contingency between physical stimuli and correct choices; for example, monkeys sometimes received reward by making a Go response to the 0% morph stimulus. Could this broken stimulus-target contingency alone account for the observed closed-loop effects? To directly test this, we designed a control experiment in which the stimulus-target contingency was broken in the same way without linking correct choices to neural activity (Fig. 4L). In this random reward task, the correct answer on each trial was assigned randomly to Go or No-go choice according to probabilities derived empirically from the closed-loop sessions for the corresponding object feature and morph level of the same monkeys. Thus, the only difference from the closed-loop experiments, hidden from the monkeys, was whether correct choices were determined by neural activity or assigned at random.

In this random reward task, we found that the neuron–choice correlation did not improve. Correlation slopes were indistinguishable between this condition and baseline in both monkeys (Fig. 4M; *t*(10) = *−*1.08, *p* = 0.31 for monkey A and *t*(10) = *−*0.39, *p* = 0.70 for monkey B, two-tailed *t*-test), despite a small increase in overall Go responses similar to that seen in closed-loop sessions. The monkeys’ psychometric functions changed (Supplementary Fig. 7) in response to the broken stimulus-target contingency, but they were not accompanied by any change in neuron-choice slope.

## Discussion

How sensory neural activity contributes to perceptual reports has been a fundamental problem in systems neuroscience. Existing approaches have relied either on measuring correlations between neural activity and perceptual choice (Nienborg et al. 2012), or on perturbing sensory areas through coarse stimulation or inactivation (Parker and Newsome 1998). We developed a third approach: a closed-loop paradigm that trains animals to use specific patterns of their own population sensory neural activity to guide decisions. We tested this paradigm in monkeys performing a change-detection task, and found that they clearly increased the correlation between their choices and population signals along the decoding axis for a visual feature in V4 (Fig. 3). This enhancement in neuron–choice correlation occurred without any detectable change in the response properties of the neural population itself (Fig. 4A-E). A computational model confirmed that the closed-loop effect was explained by adjustments in readout weights, rather than by other factors known to influence neuron–choice correlations (Fig. 4F-K). A control experiment in which only the stimulus–reward contingency was disrupted did not produce such enhancement (Fig. 4L-M).

Our results demonstrate that the correlation between sensory neural activity and perceptual reports is malleable and depends on the strategy animals use to solve tasks (Chowdhury and DeAngelis 2008; Laamerad et al. 2025; Uka and DeAngelis 2004). While choice probability (CP) was originally proposed as an index of how sensory activity contributes to perceptual decisions (Britten et al. 1996), subsequent work has revealed other dominant factors such as noise correlations (Haefner et al. 2013) and top-down feedback of choice information to sensory cortex (Bondy et al. 2018; Haefner et al. 2016; Liu et al. 2026). Our closed-loop paradigm targeted an early response window to mitigate the influence of feedback (Fig. 2A) and successfully increased neuron–choice correlation without altering neural activity patterns or noise correlations (Fig. 4). The absence of changes in neural selectivity and response statistics also provides evidence against alternative accounts, such as feature-selective attention (Bichot et al. 2005), task engagement, or additional perceptual learning (Ni et al. 2018) known to influence V4 activity, which should have been observed in our recordings if present. The magnitude of the closed-loop effect was modest (Fig. 3C), but our simulation suggests this probably approached a theoretical limit: even large changes in readout weights produce comparable shifts in neuron-choice correlation (Fig. 4G).

The successful closed-loop training suggests that the misalignment of the stimulus and choice encoding observed in perceptual tasks (Ni et al. 2018, 2022; Weiner and Ghose 2014; Zhao et al. 2020) is partly a consequence of the constraints imposed by natural behavioral training. Typical perceptual tasks train animals to maximize sensory discrimination accuracy, which may or may not require aligning stimulus and choice encoding in the recorded population as an optimization target (Pitkow et al. 2015). By contrast, our paradigm directly imposed this alignment as the training objective, probing its real performance limit. The results indicate that this limit lies above what is typically observed in perceptual tasks, supporting the view that maximizing discrimination accuracy and maximizing stimulus-choice alignment are not equivalent training objectives. Under natural training, animals may favor generic task strategies that use readout weights capable of detecting a broad range of physical stimuli (Laamerad et al. 2025; Ni et al. 2022; Srinath et al. 2026; Weiner and Ghose 2014).

Mechanistically, learning in our closed-loop task could proceed through progressive adjustment of readout weights via reinforcement learning (Law and Gold 2009), but the locus of this adjustment need not be restricted to the recorded neurons. Activity along the decoded axis must be correlated with neural signals distributed broadly across the population (Cohen and Newsome 2009; Kohn and Smith 2005; Pitkow et al. 2015) and potentially across areas (Semedo et al. 2019; Zandvakili and Kohn 2015). What monkeys may therefore have learned is to weight specific axes of latent states shared across large neuron pools. The recruitment of broad latent states is likely a general feature of closed-loop paradigms: in motor cortex, animals can be trained to control neural activity only along directions corresponding to naturally occurring latent states or dynamics (Oby et al. 2025; Sadtler et al. 2014). Our paradigm extends this logic to sensory subspaces, demarcating the limit of aligning latent states and behavioral reports. This opens further questions, for example, which sensory areas permit such alignment most easily, whether readout weights can be flexibly retargeted across different axes (see Supplementary Fig. 8), or what forms of neural information beyond linear decoding of spike rates can be linked to behavior (Panzeri et al. 2010; Quiroga and Panzeri 2009; Yang et al. 2021).

Another important factor shaping sensory readout is trial-to-trial co-fluctuations in population activity. While noise correlation can substantially reduce stimulus coding fidelity (Moreno-Bote et al. 2014; Pitkow et al. 2015), it is known that the choice encoding still tends to align with the dominant axes of noise correlations (Ni et al. 2018; Valente et al. 2021), a finding replicated in our experiment (Fig. 2E). Multiple theoretical frameworks have interpreted this alignment as evidence that shared neural fluctuations rather play a constructive role in transmitting sensory information to downstream decision circuits (Haimerl et al. 2023; Nassar et al. 2021; Panzeri et al. 2022; Srinath et al. 2025). A key observation from our closed-loop experiment is that, although stimulus–choice alignment improved following training, the choice information carried by the whole recorded population and the dominant noise axes remained stable (Fig. 3E). This dissociation might indicate that the dominant axes of population variability may act as a constraint that prevents readout weights from deviating from them substantially. This interpretation resonates with the concept of a communication subspace (Semedo et al. 2019; Srinath et al. 2021), in which inter-areal communication is constrained to a low-dimensional subspace. Under this view, closedloop training may have improved readout efficiency within the available communication subspace.

Beyond its scientific implications, the proposed closed-loop paradigm represents a conceptually novel approach to real-time neural interface, distinct from conventional brain-computer interface (BCI) designs. The dominant BCI framework, developed primarily for motor cortex applications, trains subjects to volitionally modulate their own neural activity to control an external device (Andersen et al. 2022; Motiwala et al. 2025). This framework has been extended to sensory cortices in several ways: closed-loop neurofeedback has been used to train subjects to modulate sensory or attention-related activity in visual areas and influence perceptual accuracy or contents (Amano et al. 2016; Debettencourt et al. 2015; Renton et al. 2021; Shibata et al. 2011). Recently, Iordan et al. (2024) used implicit closed-loop feedback to guide the formation of categorical representations in sensory cortex. Separately, closed-loop systems have been used to identify sensory selectivity with iterative stimulus sampling (Okazawa et al. 2015, 2017; Yamane et al. 2008). What distinguishes our paradigm from all of these approaches is that we do not aim to modulate the amplitude or pattern of recorded neural activity itself. Instead, subjects are required to change how downstream circuits read out that activity. This is a fundamentally different point of intervention in the sensory-to-decision pathway and, to our knowledge, has not been previously demonstrated.

Our approach may also offer translational insights. A major challenge in restoring sensory function to patients through neural prosthetics is that even when sensory input is successfully introduced, patients often struggle to interpret and act on these novel signals (Fine and Boynton 2015). The difficulty may partly lie in the problem of readout mechanisms linking sensory signals to action. Our closed-loop paradigm could offer a strategy to train the brain to use specific patterns of sensory activity.

## Methods

### Subjects and experimental apparatus

The experiments were performed with two adult male macaque monkeys (monkey B: *Macaca mulatta*, 10 kg, 13 years old; monkey A: *Macaca fascicularis*, 6.5 kg, 11 years old). All the experimental procedures conformed to the National Institutes of Health *Guide for the Care and Use of Laboratory Animals* and were approved by the Institutional Animal Care and Use Committee of the Center for Excellence in Brain Science and Intelligence Technology, Chinese Academy of Sciences.

Monkeys were seated in a semi-dark room with their heads stabilized using a surgically implanted head post, facing a cathode ray tube monitor (SONY Multiscan E-200; 1024 *×* 768 pixel resolution, refresh rate 75 Hz). The task and stimulus presentation were controlled with a custom MATLAB program using the Psychtoolbox toolbox (Brainard 1997). The closed-loop system ran on a separate computer using a custom program with an online spike monitoring package (cbMEX; Blackrock Microsystems, Inc., Salt Lake City, UT). The program received a TTL pulse from the task computer at stimulus onset, counted spikes within a fixed window (see below), and returned the spike count to the task computer prior to the feedback period. We confirmed that the closed-loop system successfully returned spike counts before feedback on every trial. Monkey’s gaze position was monitored at 1 kHz using an infrared eye-tracking system (EyeLink SR1000, SR-Research, Ontario).

### Change detection task

Monkeys were trained to perform a change-detection task in which they viewed two sequentially presented object stimuli and reported any change (color, shape, texture) in the second stimulus by making a saccade. In each trial, after the monkey fixated on a central fixation dot, a peripheral target dot appeared outside the average V4 receptive field (RF) location at 8 deg eccentricity (rotation angle roughly orthogonal to the average V4 RF). This location was chosen to mitigate saccade-related responses in V4 (Chandrasekaran et al. 2024; Steinmetz and Moore 2014). Following a brief delay (exponential distribution, 200 ms mean, and 50 ms standard deviation), a reference object stimulus was presented for 200 ms at the center of the average V4 RF, followed by a 300 ms blank interval, after which a test object stimulus appeared at the same location. If a change in the object was detected, monkeys had to make a saccade to the target dot within a fixed response window (180-800 ms after stimulus onset for monkey A and 200-800 ms for monkey B). If no change was detected, monkeys had to maintain fixation for 800 ms. Changes occurred in approximately 50% of trials. Correct responses were rewarded with a juice drop. The mean reaction time for choosing Go was 241 ms for monkey A, and 289 ms for monkey B.

The reference stimulus was always the same colorless, amorphous 3D object image, whereas test stimuli in Go trials differed from the reference in color, texture, or shape. To create the reference stimulus, we began with a 3D sphere model, randomly placed isotropic Gaussian bumps on its surface, and rendered a 2D image in MATLAB. Test stimuli with a feature change were created by applying one of the following five manipulations to the reference: (1) sharpness: increased the height and decreased the width of the Gaussian bumps to make them appear sharper. (2) aspect ratio: stretched the entire object shape in the horizontal direction. (3) color (red): rendered the entire surface reddish. (4) color (green): rendered the entire surface greenish. (5) texture: applied a naturalistic texture to the surface. Monkey A was tested with four manipulations (except for the color green); Monkey B was tested with all five.

To control stimulus difficulty, we created a morph continuum for each feature ranging from 0% (identical to the reference) to 100% (maximum difference). For shape and color manipulations, stimulus parameters were linearly modulated along the continuum. For texture, we linearly increased the contrast of the texture. The 100% morph of each feature was calibrated using preliminary human psychophysics such that all features yielded comparable detection thresholds (20-30%). During experiments, morph levels and their presentation probabilities were adjusted according to the monkey’s behavioral performance; we typically used 10%, 15%, 20%, 25%, 30%, 35%, 40%, and 60% levels. Approximately 50% of trials were No-go trials, in which the 0% (reference) stimulus was presented; the remaining trials were Go trials, with morph level drawn from the levels listed above.

We had two types of sessions for the change-detection task. In one type, all 4-5 features were presented to train monkeys to detect any change in object features. These sessions were used primarily for training. The other type used only one feature. These sessions served as the baseline against closed-loop sessions using the same feature. We collected 22 of these single-feature sessions for monkey A (22,419 trials) and 28 sessions for monkey B (22,475 trials). Figure 2 also used these baseline sessions.

### Closed-loop experiments

In the closed-loop task, we modified the reward contingency so that monkeys maximized reward by aligning their Go/No-go choices with their own V4 neural activity recorded online. The task structure was otherwise identical to the change-detection task. During test stimulus presentation, the closed-loop system counted the number of spikes within an 80-150 ms window from stimulus onset, projected the signals onto a neural axis defined during a preceding passive fixation task (see below), and compared the projected value against an experimenter-set threshold. If the projected neural activity exceeded the threshold, the correct response for that trial became Go, regardless of the physical stimulus presented. When we introduced this rule, we initially applied it only to 0% morph trials, so as not to confuse the monkeys in the early half of sessions (total 39 sessions for monkey A and 7 sessions for monkey B). Later, we applied the closed-loop rule to all the morph levels. Because the target neural axis was chosen to be the optimal decoding axis for the presented feature, the reward contingency was preserved on the majority of trials, but still the contingency was reversed on a substantial fraction of trials (*∼* 25 % of trials), encouraging the monkeys to learn the new rule through trial and error. No explicit cues were provided to indicate the closed-loop rule.

To identify the target neural axis for the closed-loop training, we performed a passive fixation task prior to the main task in each daily session. On each trial, 2-4 object images were presented at the center of the average V4 RF for 200 ms with a 300 ms inter-stimulus interval. We presented morph levels of 0, 20, 40, 60, 80, and 100 % for all features with *∼* 20 repetitions each, and calculated the mean firing rate of each unit for each stimulus within an 80-150 ms window from stimulus onset. Before identifying the stimulus encoding axis, we centered the neural population responses on the average responses to 0% morph stimulus and applied principal components analysis (PCA) to generate a low-dimensional latent space. The number of principal components was set to 12, based on parallel analysis (Altan et al. 2021) conducted during early data collection (Supplementary Fig. 2A).

Within this 12D space, we estimated a stimulus encoding axis using one of two methods. The first method used a linear regression of neural population response onto morph level for each feature *f* , presented *N_f_* times during the passive fixation task:

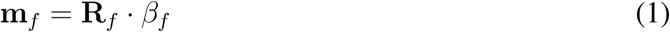

where **m***_f_* is an *N_f_* vector of morph levels, **R***_f_* is an *N_f_ ×* 12 matrix of neural population responses in the 12D latent space, and *β_f_* is a 12 *×* 1 regression coefficients, which defined the encoding axis for feature *f* . These coefficients were independently determined for each object feature. The second method, used in a part of sessions (6 sessions), estimated approximately orthogonal encoding axes for each feature simultaneously, such that they were mutually uncorrelated. Using all *N* trials of the passive fixation task across *F* features, we constructed an *N × F* matrix of morph level **M***_all_* and an *N ×* 12 matrix of neural population responses **R***_all_*. Canonical correlation analysis (CCA) was then applied to determine coefficient matrices **A** (*F × F*) and **B** (12 *× F*) that maximized the correlation between each column of **M***_all_ ·* **A** and **R***_all_ ·* **B**. The resulting matrix **BA***^−^*^1^ defines the projection from neural population activity onto the axes most correlated with the morph level of each feature.

To correct for slow drift in neural population activity, which could gradually bias the closed-loop threshold over the course of a session, we applied an online normalization procedure to the decoded neural signals. The mean and standard deviation of population activity projected on the target axis were calculated from the most recent 50 0%-morph trials, and the projected value of each trial was normalized by subtracting the mean and dividing by the standard deviation. For the first 50 trials of the closed-loop experiment, neural responses from the preceding passive fixation task were included in this normalization to provide a stable baseline estimate.

When starting the closed-loop experiment with monkey A, we initially attempted to align the latent neural spaces across sessions so that the monkey could learn a consistent rule across days (20 sessions; 21% of data). Following previous studies (Degenhart et al. 2020), we applied a Procrustes transformation to align the PCA coefficient matrix of the 12D latent neural space from day N (*Q_N_*) to that of day 1 (*Q*_1_) through a rotation matrix (*T_P_*) and a translation vector (*S_P_*), obtained by:

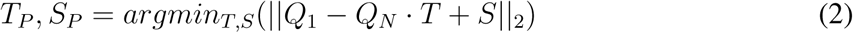

Neural activity of day N was then projected onto the aligned latent space using the transformed PCA coefficients *Q_N_ · T* + *S*. In later sessions, we found that this alignment did not improve learning stability, thus we directly used the stimulus axis defined through the passive fixation task on a session-by-session basis.

### Closed-loop experimental schedule and training

We performed baseline and closed-loop sessions over consecutive days using the same object feature within a single round of data collection (Fig. 3B). Each round began with at least 4 baseline sessions, in which monkeys performed a regular change-detection task with one object feature. We verified that behavioral performance was stable during this period. Closed-loop sessions were then conducted for at least 5 sessions (mean: 7.4). Each baseline and closed-loop session began with the passive fixation task to determine the feature axes as described above. For monkey A, a subset of baseline sessions (8 sessions) used an alternative closed-loop rule in which the target axis was set orthogonal to the object feature axis for 0% morph stimulus. This was intended to dissociate monkey behavior from a closed-loop contingency learned in the previous round. However, this manipulation did not produce substantial differences in behavior.

During closed-loop experiments, several parameters were adjusted online by the experimenters to facilitate learning. First, the threshold separating Go/No-go categories along the decoder axis was set so that Go trials comprised approximately 50% of all trials. The threshold was typically positioned around 1 standard deviation above the mean decoded value for 0% morph stimulus. Concurrently, we adjusted the proportion of 0% morph trials to ensure that monkeys continued to experience physically changing stimuli on a good fraction of trials. In some sessions, we also applied a soft threshold to provide more fine-grained feedback to monkeys. Here, both Go and No-go responses around the soft boundary were treated as correct, but the reward magnitude for the “wrong” choice was scaled according to the distance of the decoded value from the threshold. For example, if the decoded value was slightly above the threshold, a Go response yielded a large reward, whereas a No-go response yielded a substantially smaller reward. To implement this, reward magnitude was defined by a linear function of the decoded value, bounded by experimenter-defined upper and lower limits. We collected a total of 46 closed-loop sessions for monkey A (43,032 trials) and 50 sessions for monkey B (42,357 trials).

### Control experiments

#### Random reward task

To test whether a broken stimulus-reward contingency alone could have induced the changes observed in closed-loop experiments, we designed a random-reward task. In this task, the contingency between physical stimuli and trial outcomes was altered in the same manner as in closed-loop sessions, but reward delivery was determined randomly rather than by online neural signals. For each morph level, we calculated the probability that a Go response was rewarded in the corresponding closed-loop sessions of the same monkey, and then used this probability to randomly reward a Go response. As a result, some 0% morph trials were rewarded for Go responses, and some positive morph trials were rewarded for No-go responses, mirroring the statistical structure of the closed-loop experiments. As in the closed-loop rounds, both monkeys first completed five consecutive baseline change-detection sessions before performing the random-reward task for eight sessions. For analysis, we included the last four baseline sessions and all random-reward sessions. We collected a total of 8 random-reward sessions for monkey A (6,333 trials) and 8 sessions for monkey B (5,411 trials).

#### Closed-loop axis switching task

In this task (Supplementary Fig. 8), we tested whether monkeys could modulate neuron-choice correlations selectively between two different target neural axes associated with two object features (e.g., sharpness and color). To prevent external stimuli from becoming a cue to the target axis, we created a morph continuum along which both features co-varied simultaneously with the same morph levels. In baseline conditions, monkeys were required to detect any change that deviated from the reference stimulus. To ensure that monkeys detected both features, catch trials with a change in one feature alone were introduced in *∼*8 % of trials. During the closed-loop sessions, we defined the decoder axis using only one of the two features and used this axis to determine the correct answer on each trial. We first collected baseline sessions for at least four sessions for each monkey, and then proceeded to closed-loop training on each of the two features sequentially (Supplementary Fig. 8A). During closed-loop sessions, the catch trials with a single-feature change were randomly rewarded with 80% probability so as not to interfere with the closed-loop training. All other parameters and procedures for the closed-loop were the same as those in the main experiment.

### Electrophysiological recording

We implanted a 96-channel Utah array (electrode length 1 mm; spacing 0.4 mm; Blackrock Microsystems) in area V4 of each monkey for chronic recordings. The implant location was planned based on a T1 MRI image and confirmed during the surgery by identifying the lunate and superior temporal sulcus. During recordings, neural signals were sampled at 30,000 Hz (high-pass filtered at 250 Hz), and waveforms exceeding a fixed threshold were detected as multi-unit activity (MUA) for closed-loop experiments, consistent with the approach used in previous closed-loop studies (Oby et al. 2025; Sadtler et al. 2014). The spike detection threshold was set to 3.0 or 3.5 times the root-mean-square (RMS) of the continuous signal for monkey A and 3.5 RMS for monkey B. Because the closed-loop experiments were performed using online MUA, the same MUA was used for analyses of both closed-loop and baseline sessions to ensure direct comparability (Figs. 3, 4). We used all channels that showed clear visual responses (*n* = 90 *−* 96). We assessed the quality of recording by performing preliminary analysis of stimulus classification and excluded two rounds of data collection from monkey A due to poor quality. For analyses that do not depend on the closed-loop paradigm (Fig. 2), we also performed spike sorting offline with Offline Sorter (Plexon, Dallas, TX 75206, USA) on a subset of the sessions to confirm consistency of results (Supplementary Fig. 9). Approximately 40-150 single and multi-units (average 100) were isolated per session.

Prior to the main experiments, we mapped the receptive fields (RFs) of V4 neurons to determine the stimulus presentation location. While monkeys passively fixated at the center of the screen, a sequence of Gabor stimuli (4 orientations, 1.5 cycles/deg) was presented at random locations on a grid with 1*^◦^* spacing. We calculated the mean firing rate of each channel at each location and fitted an isotropic 2D Gaussian kernel to estimate its RF center and size (*∼*68 % of channels successfully fit). The RF centers were then averaged across channels to determine the stimulus location ([4.2*^◦^, −*4.1*^◦^*] in Cartesian coordinate, size 5.3*^◦^*, for monkey A; [5.0*^◦^, −*2.0*^◦^*], size 5.3*^◦^* for monkey B).

### Data analysis

#### Psychometric curve fitting

Monkeys’ psychometric functions were calculated as the probability of a Go response as a function of stimulus strength (morph level) and were fitted using a generalized linear model (GLM) with a logistic link:

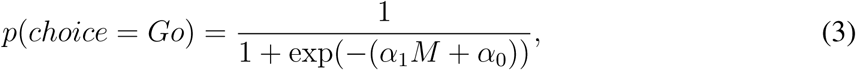

where *M* is the morph level (ranging from 0 to 1) of the test stimuli, *α*_1_ and *α*_0_ are the slope and bias parameters, respectively.

#### Stimulus and choice decoding analysis

To confirm that the recorded population linearly represented the object feature (Fig. 2C), we quantified how much variance in morph level could be explained by a linear decoder constructed in the 12-dimensional latent space (see above) using data collected during passive fixation. In each session, we decoded morph levels using a linear decoder with 5-fold cross-validation. In each fold, the training set was used to estimate a morph decoding axis, and the held-out test trials were projected onto this axis. This procedure was repeated for all five folds, and the resulting projected values were concatenated to obtain cross-validated morph level predictions for all trials. The variance explained (*R*^2^) was computed as follows:

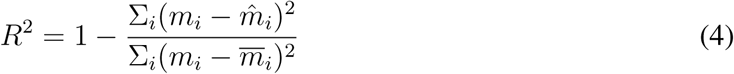

where *m_i_* denotes the actual value of each morph level *i*, *m*^*_i_* denotes the predicted morph level averaged across trials, and *m* is the mean of the morph levels.

To identify a neural axis optimally decoding the monkeys’ choices, we performed a logistic regression on neural population activity after removing the stimulus-dependent component (Fig. 2E-F). Trials with morph levels yielding *>*80% Go responses were excluded, as they were expected to carry minimal choice-related variance. Beginning from the 12D latent activity in PC space, we first subtracted the trial-averaged responses to each morph level. Logistic regression was then applied to the residual activity using leave-one-out or 5-fold cross-validation; in each fold, the training trials were used to estimate a choice decoding axis, and the held-out test trial was projected onto this axis. This procedure was repeated across all trials, and the projected values were concatenated to obtain cross-validated projections for all trials. Finally, we computed the area under the receiver-operating characteristic curve (AUC) from these cross-validated projections to quantify the accuracy of classifying Go and No-go response trials.

Choice decoding accuracy along the stimulus encoding axis and the dominant axes of neural variability (Fig. 2E-F) was computed in a similar way, after removing stimulus-dependent signals. For the stimulus axis, we projected the 12D latent signals onto this axis, subtracted trial-averaged responses to each morph level, and computed the AUC between Go and No-go response trials. The dominant axes of neural variability were defined as the top principal components (PCs) of the 12D latent space. For the first PC, the same procedure was used to subtract the stimulus-dependent signals and calculate choice decoding accuracy. For higher PCs (up to PC *k*), we estimated the optimal axis decoding choice within the subspace spanning PC 1 to *k* by applying the logistic regression described above, following the same cross-validation procedure. In Fig. 3E, we performed the same analyses but using 0% morph trials only.

Stimulus decoding performance was measured as the classification accuracy of 0% versus non-0% (Go) stimuli based on neural responses projected onto the stimulus, choice, and PC axes (Figs. 2E-F, 3E). For one-dimensional axes, accuracy was computed directly as the AUC of the projected distributions. For higher-dimensional PC spaces, we used logistic regression with leave-one-out cross-validation, as described above.

All analyses computed the decoding performance within each session and averaged it across sessions.

#### Neuron-choice correlation plot

To quantify the strength of the neuron–choice relationship, we computed the probability of a Go response as a function of neural activity projected onto the decoder axis (Figs. 3C, 4M). For each session, only 0% morph trials were used to avoid confounding stimulus information. GLM with a linear link function was used to fit monkey choices on individual trials using the projected neural signals as the predictor:

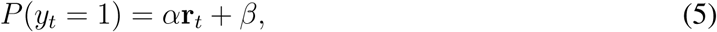

where *y_t_ ∈ {*0, 1*}* is the binary Go/No-Go choice on trial *t*, **r***_t_* is neural signal along the target axis, and *α* and *β* are the fitted slope and intercept, respectively. We chose a linear link function because the observed relationship between projected activity and Go probability appeared approximately linear (Fig. 3C) within the restricted range of neural activity evoked by 0% morph stimulus. To enable comparison of slopes across sessions, projected neural signals were z-scored within each session before fitting. For visualization in Figs. 3C, 5B, we binned trials into seven groups according to their projected signal values. As a control, we also estimated the neuron-choice slope using both 0% and positive morph trials (Supplementary Fig. 3A). We selected positive morph levels that fell in the same range on the decoder axis as the zero morph trials (mean *±* 3 s.d.) for this analysis, as Go probability saturates at higher morph levels and introduces a ceiling effect.

#### Quantification of population stimulus selectivity

To assess stimulus selectivity (Fig. 4A), we performed 5-fold cross-validation, in which we re-estimated the stimulus encoding axis on the training trials (Eq. 1) in each fold and projected the neural activity of the held-out trials onto this axis. Pooling the projected values across all 5 folds, we computed AUC between 0% morph and each positive morph level, yielding a classification accuracy (*A_m_*) for each morph level *m*. These values were fit with a logistic function to estimate a neurometric threshold:

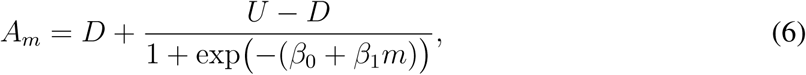

where *U* and *D* define the upper and lower limits of the function *A_m_*, *β*_0_ is a bias term, and *β*_1_ is a neurometric slope. We fitted these parameters and compared neurometric slopes *β*_1_ between baseline and closed-loop sessions using a two-sample *t*-test.

#### Noise correlation structure

We tested whether noise correlations changed following closed-loop training (Fig. 4C). Noise correlations (spike count correlations) were calculated using 0% morph trials to exclude stimulus-dependent correlations. We counted spikes within a 80-150 ms window after stimulus onset, and calculated Pearson’s correlations across all valid channel pairs. The resulting pairwise correlations were averaged within each session.

We also checked the relationship between neuronal tuning and noise correlation (Haefner et al. 2013). Since most units exhibited monotonic tuning to morph level (Fig. 4E), we quantified tuning strength for each unit using linear regression:

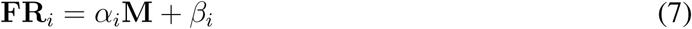

where **FR***_i_* is the vector of trial-average firing rates across morph levels for unit *i*, *M* is the corresponding vector of morph levels, and *α_i_* and *β_i_* are the regression slope and intercept, respectively. The tuning difference between units *j* and *k* was then defined as the absolute difference in slopes:

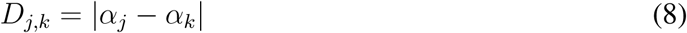

Figure 4E shows the relationship between this tuning difference and pairwise noise correlation (*ρ_j,k_*). Their relationship was quantified using another linear regression:

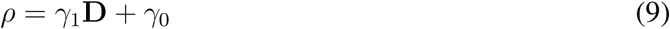

where **D** and *ρ* are tuning differences and noise correlations concatenated across all unit pairs within a session.

### Neuron pool model

#### Model structure

We built a feedforward model consisting of V4-like neural populations to simulate how different factors could affect neuron–choice correlations. The model comprises a pool of neurons (*n* = 10, 000) responding to sensory stimuli and a fixed readout from the pool that determines model choice. Firing rate of each neuron *k* on trial *i*, *FR_k,i_*, is modeled as a linear tuning to morph level (*M_i_*):

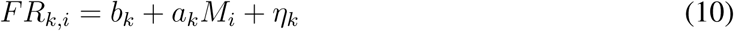

where *b_k_* is the neuron’s firing rate for 0% morph stimulus, *a_k_* is its tuning slope, and *η* is the noise level. Importantly, these parameters were sampled from the distribution of actual neural responses from all units from all sessions. For *a_k_* and *b_k_*, we performed the linear regression defined in Eq. 7 for each real unit to obtain the parameter distributions and sampled the values from their joint distribution through kernel density estimation. The noise level (*η_k_*) was quantified as the Fano factor calculated using 80-150 ms window from stimulus onset, and the values for the simulation were sampled from the distribution obtained from the actual data.

We then determined the correlation structure of these neurons, incorporating the pairwise correlation distribution of actual units. The following linear function was used to approximate the pattern of noise correlation as a function of tuning similarity (Law and Gold 2009):

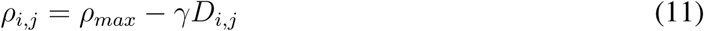

where *ρ_i,j_* is noise correlation between unit *i* and *j*, *D_i,j_* is tuning difference between the units (Eq. 8), *ρ_max_* is the maximum correlation, and *γ* is the slope defining the decay of correlation with tuning difference. The latter two parameters, *ρ_max_* and *γ*, were obtained by fitting actual data aggregated across sessions. Then, we generated the noise correlation matrix for the 10,000 simulated neurons.

Having defined the mean and correlations of simulated neural population responses, we sampled 1,000 simulated trials, similar to typical trial counts in a session. Following the method by Law et al. (2009), a 1, 000 *×* 10, 000 matrix of Gaussian noise with the correlation defined in Eq. 11 was created using *mvnrnd* function in Matlab, and then converted to firing rate based on the tuning slope for morph level, *a* (Eq. 10), the base firing rate for 0% morph stimulus, *b*, and Fano factor for each simulated neuron. Because this conversion procedure affects noise correlations, we determined *ρ_max_* in Eq. 11 such that the distribution of noise correlations approximates the data after this conversion (Law and Gold 2009). Because this simulation is stochastic, we generated five independent firing-rate matrices and presented the standard deviation of results across these samples as error bars in Fig. 4G-J to demonstrate the model stability.

From simulated firing rates, model choice was determined based on a linear readout defined by a 10, 000 *×* 1 weight vector. If the readout value exceeded a threshold (*B_dec_*), the model chose Go on that trial. A weight vector for the baseline condition was created by adding random noise to the optimal stimulus decoding weight to mimic the dissociation of stimulus and choice encoding in actual data (Fig. 4F). The optimal weight (*w_optim_*) can be determined by the following formula (Pitkow et al. 2015):

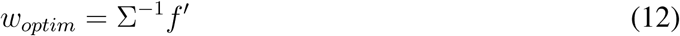

where *f ^′^* is the derivative of neuron tunings (i.e., tuning slope *a_k_* in Eq. 10) and Σ is the neuron covariance matrix. This optimal weight was then mixed with a random noise weight (*w_rand_*) drawn from a Gaussian distribution after normalizing each of them:

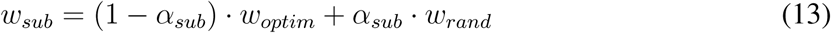

where *α_sub_* controls the degree of suboptimality. After *w_sub_* was normalized, the decoded value was calculated as **FR***^T^ w_sub_*. A Gaussian decision noise (SD, *σ_dec_*) was further added to this decoded value before it was compared against the decision threshold (*B_dec_*).

We then simulated neural recordings by randomly selecting 100 neurons from the 10,000 simulated neuron pool. Within these selected neurons, we estimated the stimulus encoding axis as we did for the actual data and calculated the neuron-choice slope as in Fig. 3C. Prior to this calculation, we added Gaussian noise to the simulated recording data (SD, *σ_record_*) to mimic the noise in neural recordings.

To summarize, the model had five parameters to generate simulated V4 populations, which were derived from the value distributions of actual neural data: tuning slope for morph level (*a_k_* in Eq. 10), baseline firing rate for 0% morph stimulus (*b_k_* in Eq. 10), variability of firing rate (*η_k_* in Eq. 10), maximum noise correlation (*ρ_max_* in Eq. 11), and the change in correlation as a function of tuning difference (*γ* in Eq. 11). The model then had four additional free parameters that could not be estimated from neuron data and should have been set by the experimenters: suboptimality of readout weights for baseline condition (*α_sub_* in Eq. 13), decision threshold (*B_dec_*), decision noise (*σ_dec_*), and recording noise (*σ_record_*).

#### Adjustment of model parameters for baseline and closed-loop conditions

To first approximate behavioral psychometric functions and neuron-choice correlations in the baseline condition, we manually adjusted the free parameters, while the parameters related to neural responses were held fixed to those derived from actual neural data. By setting *α_sub_* = 0.89, *σ_dec_* = 11, *σ_record_*= 38, and *B_dec_* such that the probability of Go choices for 0% morph was 0.16, the model could generate patterns similar to the baseline condition in monkey B (Supplementary Fig. 6B).

We then tested whether increasing the readout weights for the recorded neurons could reproduce the increase in neuron-choice correlation in the closed-loop condition. To implement this manipulation, we created another vector of readout weight, *w_rec_*, which was set to 1*/*10 for the 100 recorded neurons in the simulation and set to 0 for the rest of the simulated neurons (the norm of *w_rec_* was 1). Thus, this axis was optimally aligned to decode the signals of the recorded neurons. The readout weights were then generated by mixing this *w_rec_* and *w_sub_* in Eq. 13 as:

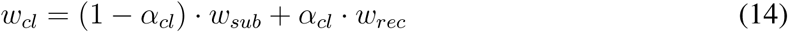

where *α_cl_* is the strength of alignment with the axis of recorded neurons. Aside from the readout weights, no other parameters were modified. The resulting neuron-choice correlation slopes are shown in Fig. 4G.

To test whether changing other parameters could reproduce the closed loop effects, we varied the following parameters controlling the property of neural responses (Fig. 4H-K): (1) tuning slope *a*, (2) the maximum noise correlation *ρ*, (3) the slope *γ* of noise correlation as a function of tuning difference. As in the baseline model, we generated five independent simulated population responses for each set of parameters to confirm the consistency of model outcomes. We also changed the decision threshold (*B_dec_*) to test if the change in the overall rate of a Go response could affect the neuron-choice correlation (Supplementary Fig. 6C).

## Acknowledgments

We thank Yong Gu and Qianli Yang for the comments on earlier versions of the manuscript. We thank Mengya Xu for technical support. This work was supported by the Strategic Priority Research Program of the Chinese Academy of Sciences (XDB1010202), the National Science and Technology Innovation 2030 Major Program (Grant No. 2021ZD0203703), National Natural Science Foundation of China (Grant No. 32371077 and No. W2432019), and the Shanghai Municipal Science and Technology Major Project (Grant No. 2019SHZDZX02).

## Competing interests

The authors declare no competing interests.

## Data and code availability

Data and code used in this study will be made publicly available upon official publication.

## Supplementary figures

**Figure S1:**
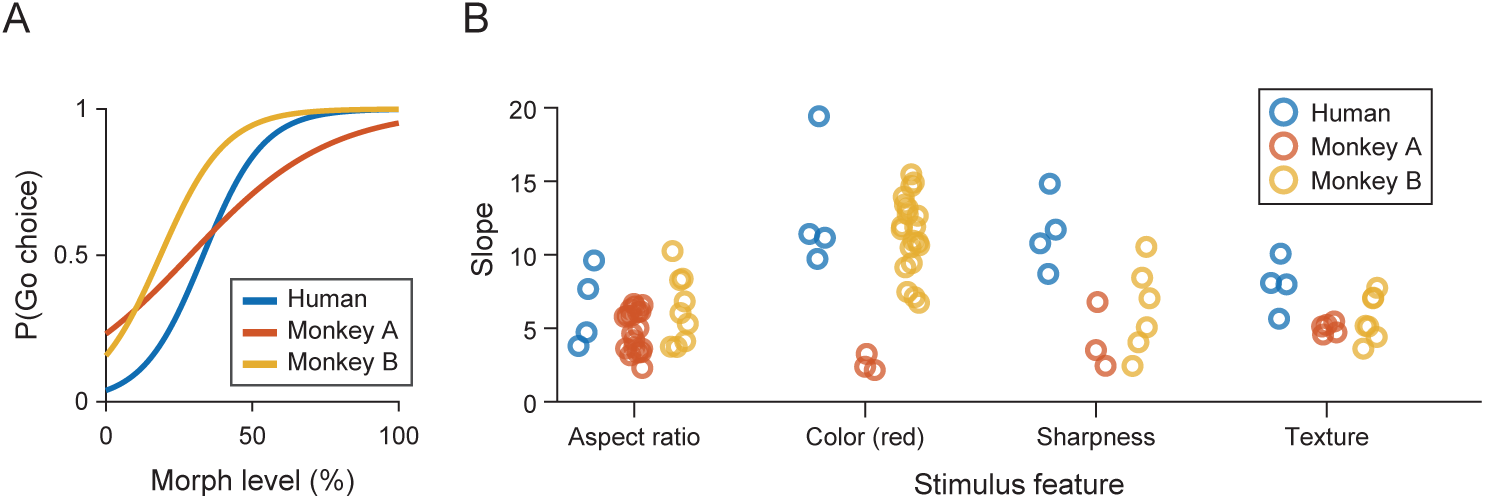
Monkey behavioral performance in the change-detection task. (**A**) Psychometric curve averaged across all object features for two monkeys and a human subject. Human data were collected using the same task structure at a stimulus eccentricity comparable to that used in monkey experiments (stimulus position: h=4.0°, v=4.0°). (**B**) Psychometric slopes (α_1_ in Eq. 3) for individual sessions. Each data point represents one session.

**Figure S2:**
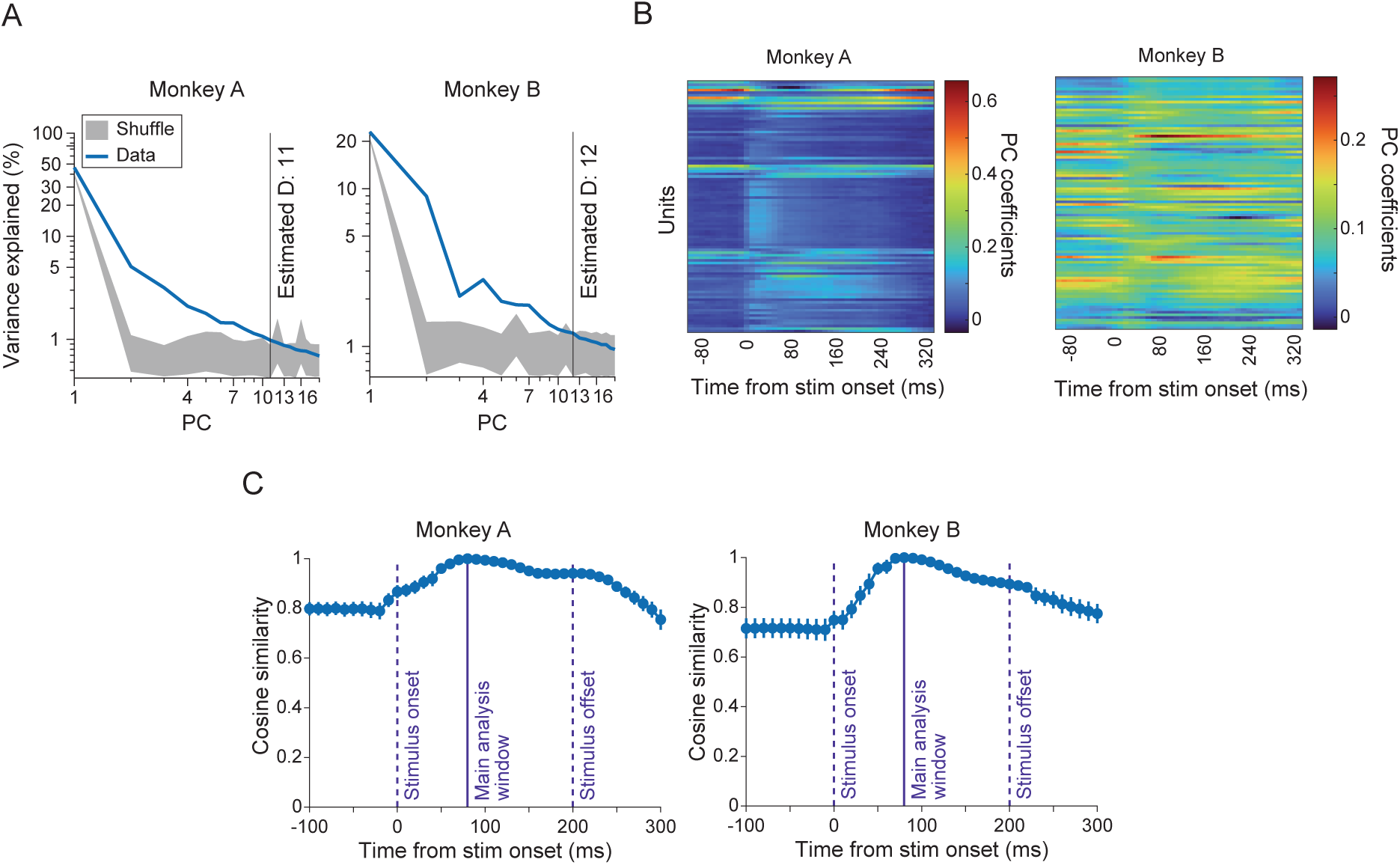
Properties of noise correlation structures in the recorded populations. (**A**) To determine the dimensionality of latent states, we performed a parallel analysis (Altan et al. 2021) in a passive fixation session conducted at the beginning of electrophysiological recording. The explained variance of each principal component (PC; blue line) was compared against a null distribution (gray) generated by shuffling unit labels. The latent state dimensionality was defined as the number of PCs whose explained variance significantly exceeded the null. (**B, C**) The dominant noise-correlation axis (PC1) remained stable across task events. We performed PCA using a 70-ms sliding window and plotted PC coefficients across channels (B). The main analysis used a window that started 80 ms after stimulus onset. The cosine similarity of PC coefficients between this reference window and all other windows remained high, including periods before and after stimulus presentation (C).

**Figure S3:**
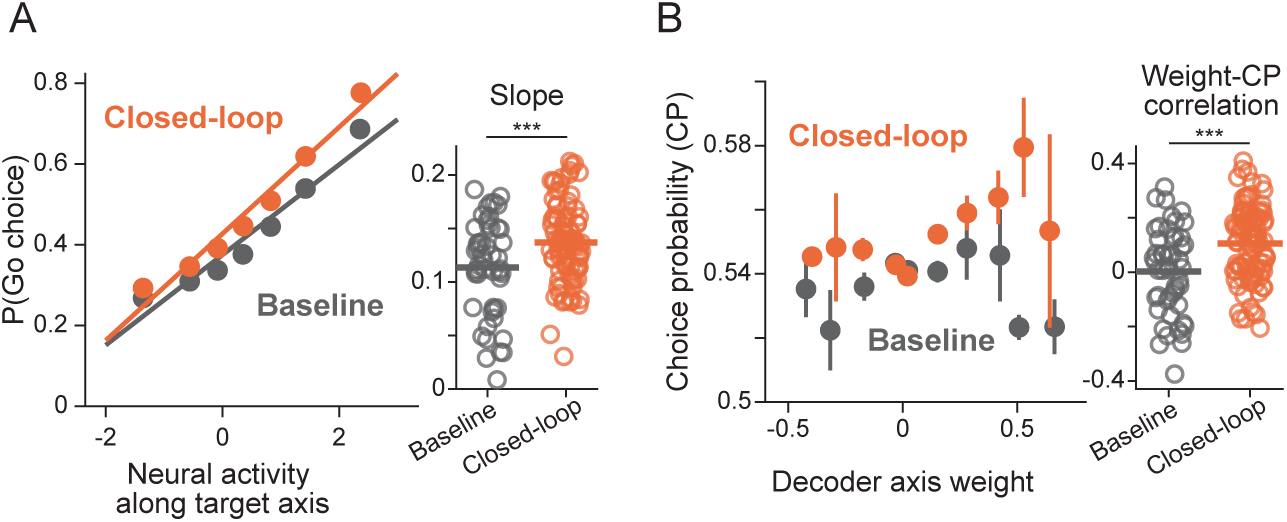
Additional analyses for the closed-loop sessions. (**A**) Neuron-choice correlation computed including trials with non-zero morph levels (< 3 SD of the mean stimulus axis value of zero morph stimulus; mostly 5-60% morph). The increase in slope for closed-loop sessions remained significant. If we include even higher morph levels, Go choice probability approaches 100% and the slopes decrease due to a ceiling effect. *** p < 0.001. (**B**) Choice probability (CP) calculated for individual units showed steeper correlations with the weights assigned to these units for the closed-loop axis. The units were sorted and binned according to the decoder weights, and their average CPs were computed.

**Figure S4:**
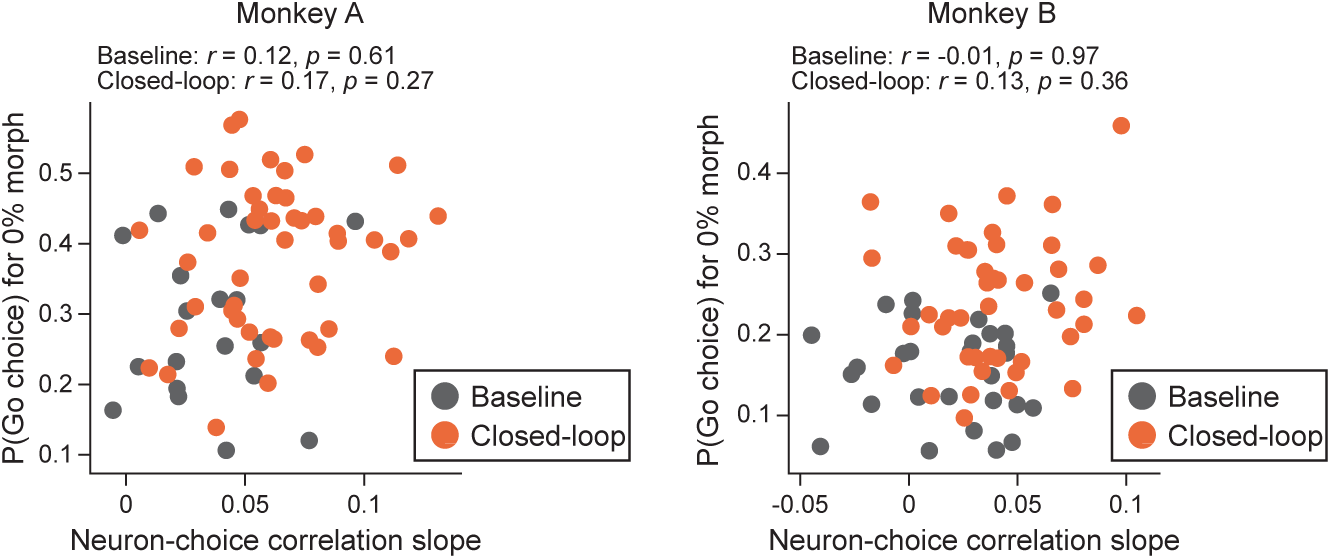
The increase in Go choice rate does not account for the closed-loop effect. Closed-loop training increased the Go choice rate alongside the neuron-choice correlation slope (Fig. 3C). To assess whether these changes were related, we plotted Go rate against slope for individual sessions (dots). No significant correlation was found in either baseline or closed-loop sessions, suggesting that the two effects were independent.

**Figure S5:**
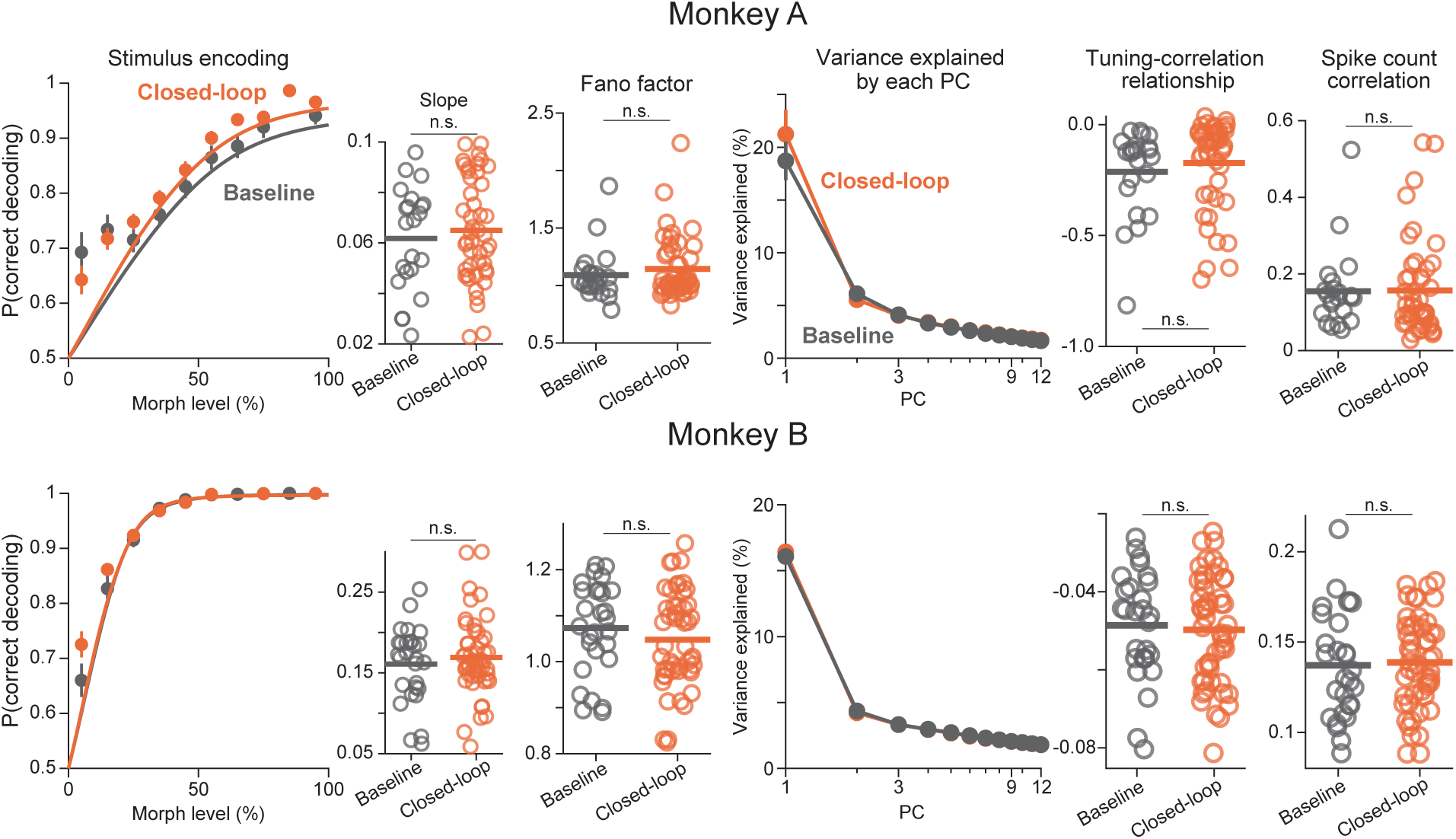
Individual monkey results for the stability of neural response properties across baseline and closed-loop sessions. Individual monkey data corresponding to Fig. 4A-E. In both monkeys, all examined properties of the neural population responses were statistically indistinguishable between baseline and closed-loop sessions.

**Figure S6:**
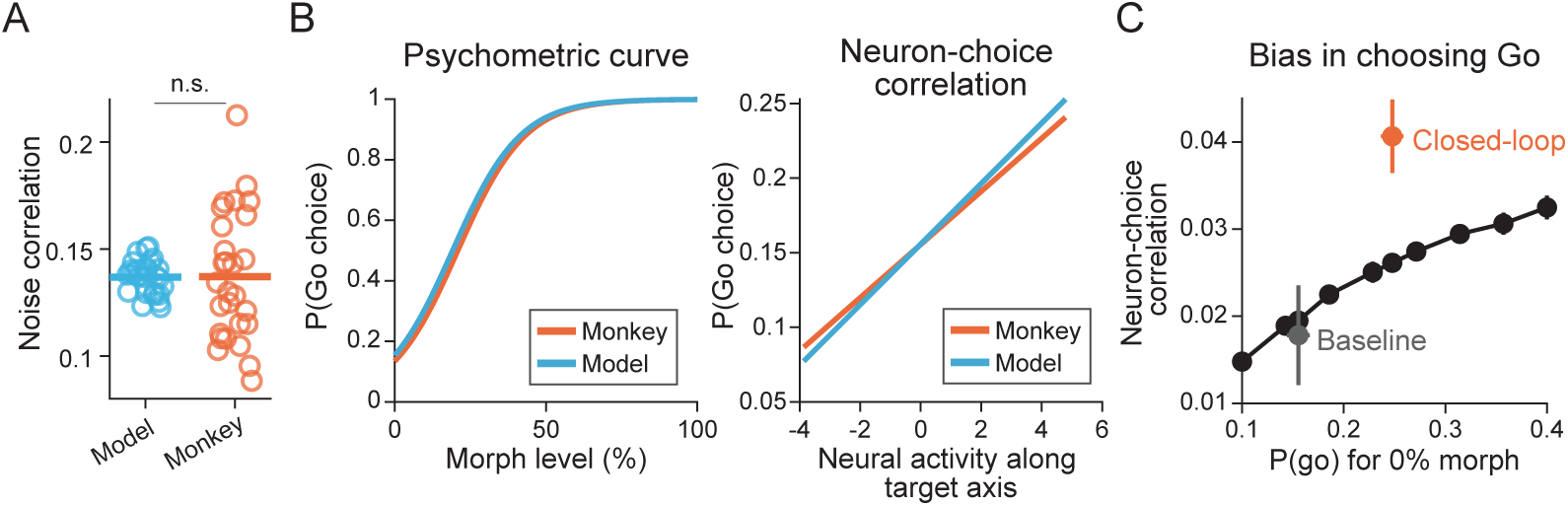
Additional analyses of the neuron pool model simulating the closed-loop effect. (**A**) To construct the neuron pool model (Fig. 4F), we built a sensory neuron pool based on the tuning strength and noise-correlation structure of the recorded neurons. After generating simulated spike counts, we confirmed that the noise correlations of the simulated data matched those of the actual neural data. Data points for the model represent individual simulations. Data points for the data are from baseline sessions of monkey B. (**B**) By adjusting model parameters, including readout weights, decision threshold, decision noise, and recording noise (see Methods), the model reproduced both monkey behavior (left) and neuron-choice correlation slope (right) in the baseline sessions. Data are averaged across baseline sessions from monkey B. Model parameters were fit to monkey B, but the model could fit monkey A data equally well. (**C**) Changing the decision threshold could not replicate the neuron choice correlation change from baseline (gray) to the closed-loop condition (orange).

**Figure S7:**
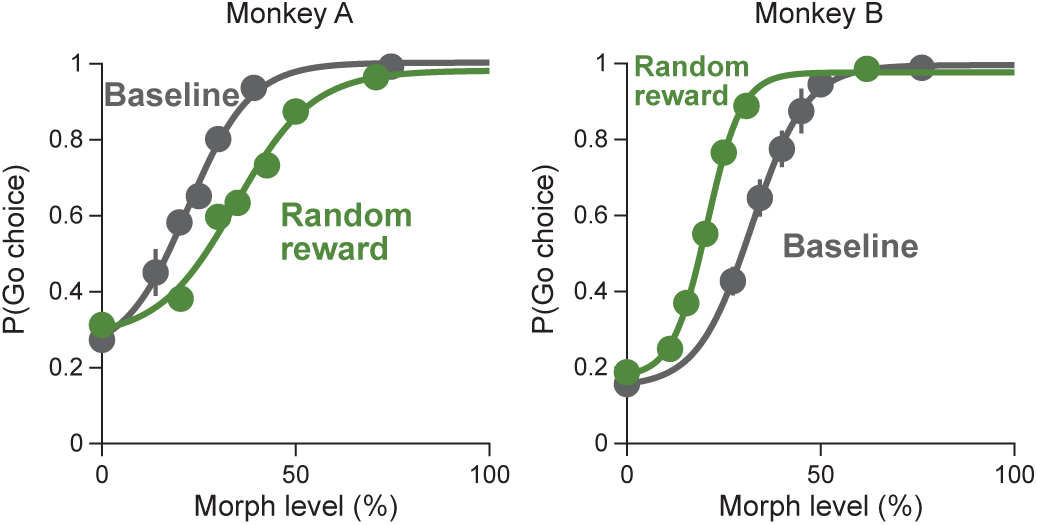
Psychometric functions in the random-reward task. The random-reward design (Fig. 4L) produced different behavioral effects in the two monkeys. Monkey A showed shallower psychometric slopes, while monkey B showed higher Go rates. This likely reflects different strategic responses to partial reward randomization; monkey A became more stochastic, whereas monkey B increased Go responses, mirroring the behavior in closed-loop sessions. Neither reaction, however, was accompanied by an increase in the neuron-choice correlation slope (Fig. 4M).

**Figure S8:**
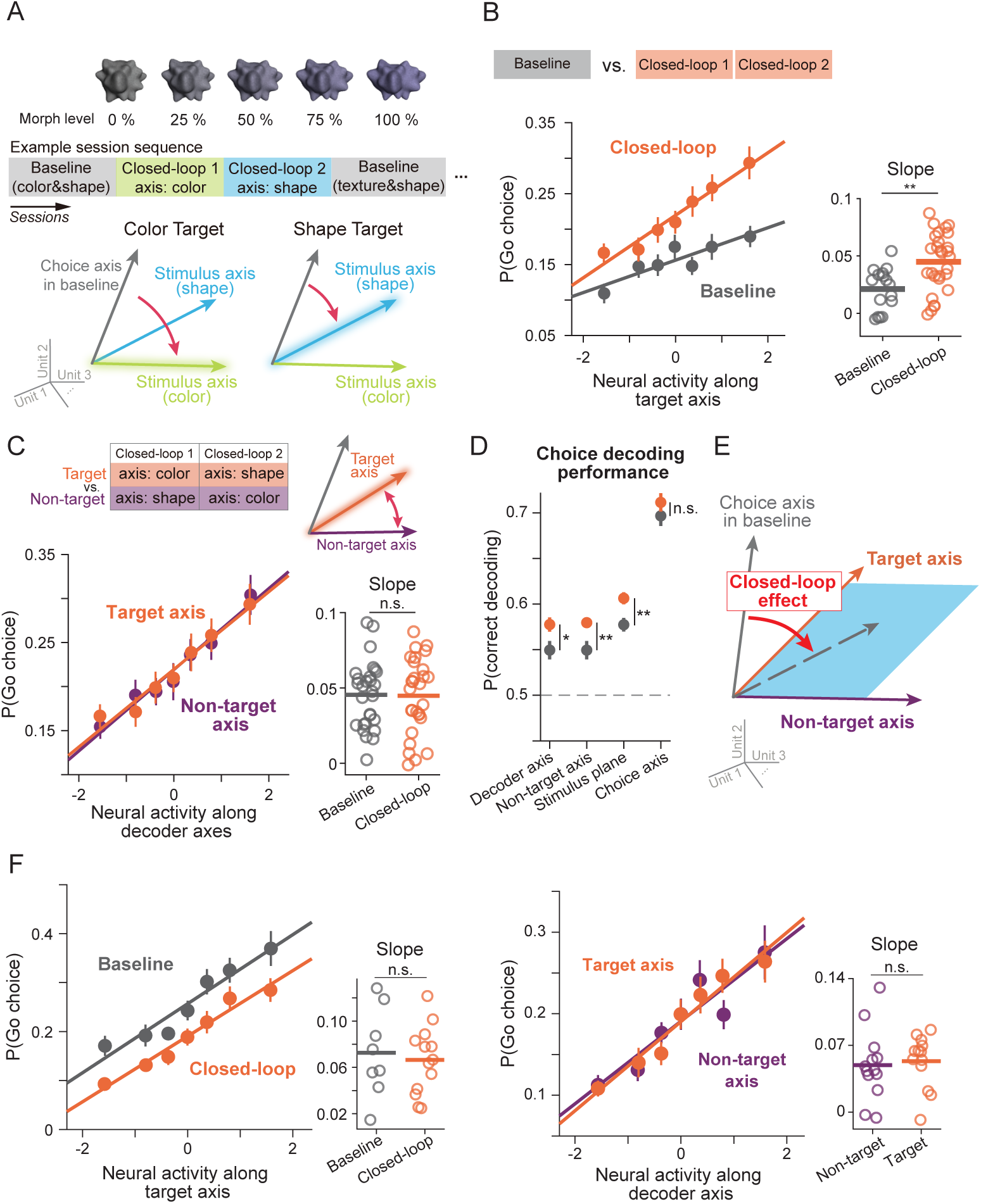
Closed-loop training induces the alignment to a feature-generic plane. (**A**) We performed an additional preliminary experiment in which the target axis for the closed-loop was alternated between two feature axes across sessions. To ensure external stimuli did not cue the target axis, stimuli were drawn from a morph continuum along which color and shape co-varied, such that both features shared the same morph level at all times (top). (**B**) Neuron–choice correlation increased during closed-loop sessions compared with baseline sessions in monkey B (p = 0.002, two-tailed t-test). (**C**) Despite an overall increase from baseline, the slopes along the target and non-target axes were statistically indistinguishable (p = 0.90, two-tailed t-test), suggesting that the monkey aligned its choices with a feature-generic axis. (**D**) Correspondingly, choice decoding performance improved along both the target axis and nontarget axes. Furthermore, the improvement looked cleanest along the axis that best decoded the choice within the 2D subspace spanned by the two feature axes (“Stimulus plane”). The overall choice information in the full latent space did not significantly change (“Choice axis”). (**E**) Schematic. Regardless of which stimulus axis was designated as the target, closed-loop training shifted neuron-choice correlations toward the stimulus encoding plane defined by the two feature axes in this task. (**F**) Monkey A did not show an increase in neuron-choice correlation slope in this task. Consistent with this, we confirmed that neuron-choice correlation slopes were indistinguishable between the target and non-target features.

**Figure S9:**
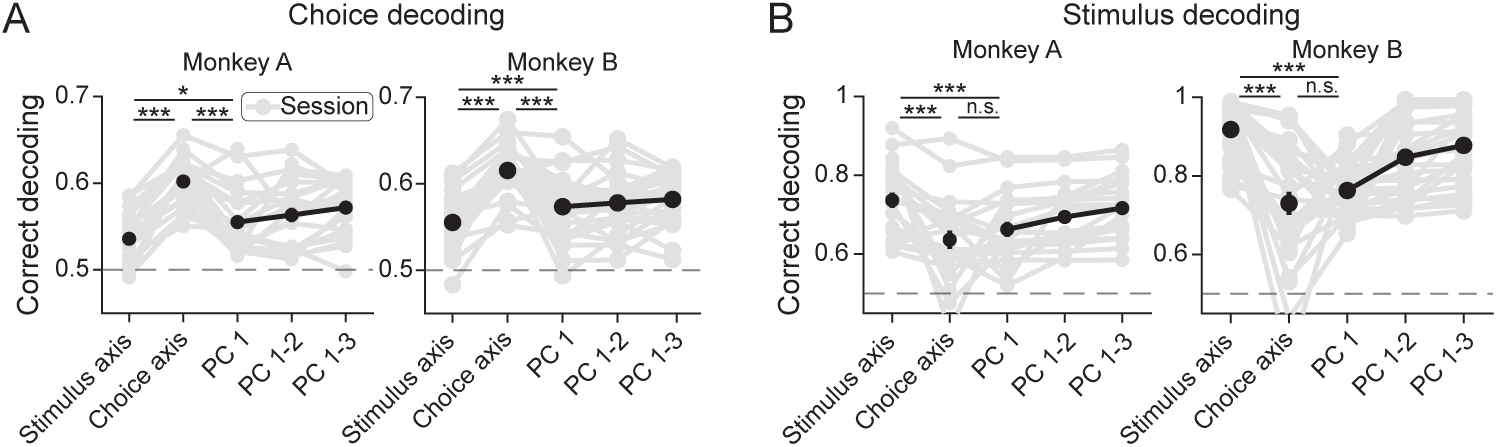
Stimulus and choice misalignment in the baseline condition is unaffected by spike sorting. Experiments used multi-unit activity (MUA) detected online (see Methods). We also performed offline sorting of the baseline sessions and confirmed the consistency of the results.

## Notes

### Competing Interest Statement

The authors have declared no competing interest.

